# Repeated strand invasion and extensive branch migration are hallmarks of meiotic recombination

**DOI:** 10.1101/2021.04.20.440609

**Authors:** Jasvinder S. Ahuja, Catherine S. Harvey, David L. Wheeler, Michael Lichten

## Abstract

Currently favored models for meiotic recombination posit that both noncrossover and crossover recombination are initiated by DNA double strand breaks but form by different mechanisms, noncrossovers by synthesis dependent strand annealing, and crossovers by formation and resolution of double Holliday junctions centered around the break. This dual mechanism hypothesis predicts different hybrid DNA patterns in noncrossover and crossover recombinants. We show that these predictions are not upheld, by mapping with unprecedented resolution, parental strand contributions to recombinants at a model locus. Instead, break repair in both noncrossovers and crossovers involves synthesis-dependent strand annealing, often with multiple rounds of strand invasion. Crossover-specific double Holliday junction formation occurs via processes that involve branch migration as an integral feature and that can be separated from break repair itself. These findings reveal meiotic recombination to be a highly dynamic process and prompt a new view of the relationship between crossover and noncrossover recombination.

## INTRODUCTION

Meiosis produces haploid gametes from a replicated diploid genome by two rounds of chromosome segregation. The first (meiosis I) involves separation of homologous chromosomes of different parental origin (homologs). Homologous recombination is crucial to this process, by promoting homolog recognition and pairing (Weiner and Kleckner, 1994) and by connecting homologs with crossovers (COs) that ensure accurate meiosis I segregation (Buonomo et al., 2000; Hassold and Hunt, 2001; Wang et al., 2017). Meiotic recombination is initiated by DNA double strand breaks (DSBs) that form at high levels (de Massy, 2013) by the conserved Spo11 protein and associated accessory proteins (Yadav and Claeys Bouuaert, 2021). DSBs are resected 5’ to 3’ to form ∼1kb ssDNA overhangs, which then invade an intact template and form a displacement loop (D-loop), followed by extension by DNA synthesis of the free 3’ end in the D-loop.

The currently favored model for meiotic recombination (Figure 1A), here called the dual-mechanism model, derives mainly from studies in budding yeast. It suggests that noncrossover (NCO) and CO recombination diverge soon after D-loops form (Allers and Lichten, 2001a). NCOs are proposed to be formed by synthesis-dependent strand annealing (SDSA), where helicase-mediated displacement frees the extended invading strand to anneal with the other resected DSB end (Allers and Lichten, 2001a; McMahill et al., 2007; Nassif et al., 1994). In contrast, COs are thought to form by a process called, for historical reasons, double-strand break repair (DSBR; Szostak et al., 1983). During DSBR, inherently unstable D-loops are stabilized by the conserved ZMM protein ensemble and converted to more stable 3-armed structures called single end intermediates (SEI; Hunter and Kleckner, 2001), which captures the second DSB end to form a four-arm double Holliday junction (dHJ) intermediate (Börner et al., 2004; Hunter and Kleckner, 2001; Lynn et al., 2007; Pyatnitskaya et al., 2019; Schwacha and Kleckner, 1995). The ZMM proteins include complexes (Zip2-Zip4-Spo16 and Msh4-Msh5) that bind branched DNA (De Muyt et al., 2018; Snowden et al., 2004), the Mer3/Hfm1 3’-5’ DNA helicase, thought to extend D-loops (Mazina et al., 2004), and E3 ligases (Zip3 in budding yeast, RNF212 and HEI10 in others) that promote ZMM protein accumulation at CO-designated sites and degradation elsewhere (Ahuja et al., 2017; Cheng et al., 2006; Qiao et al., 2014; Rao et al., 2017). Most ZMM-dependent dHJs are resolved as COs by the meiosis-specific Mutl*γ* (Mlh1/Mlh3)-Exo1 resolvase (Allers and Lichten, 2001a; Argueso et al., 2004; Cannavo et al., 2020; Keelagher et al., 2011; Kulkarni et al., 2020; Zakharyevich et al., 2010; Zakharyevich et al., 2012). A minor fraction (∼15-20%) are thought to be resolved as both COs and NCOs by nucleases active in both mitosis and meiosis (De Muyt et al., 2012; Holloway et al., 2008; Kaur et al., 2015; Kohl and Sekelsky, 2013; Zakharyevich et al., 2012). In addition, ∼20% of meiotic recombination intermediates contain 3 or 4 chromosomes; these can be produced when the two DSB ends invade different chromatids or by repair of two independent DSBs (Figure S1A; Oh et al., 2007). Finally, while most NCOs are thought to be formed by SDSA, a process called dissolution (Figure S1B), where helicases and topisomerases take apart dHJs without cleavage, has been proposed as an alternate (Dayani et al., 2011; Gilbertson and Stahl, 1996; Nasmyth, 1982; Wu and Hickson, 2003).

**Figure 1.**
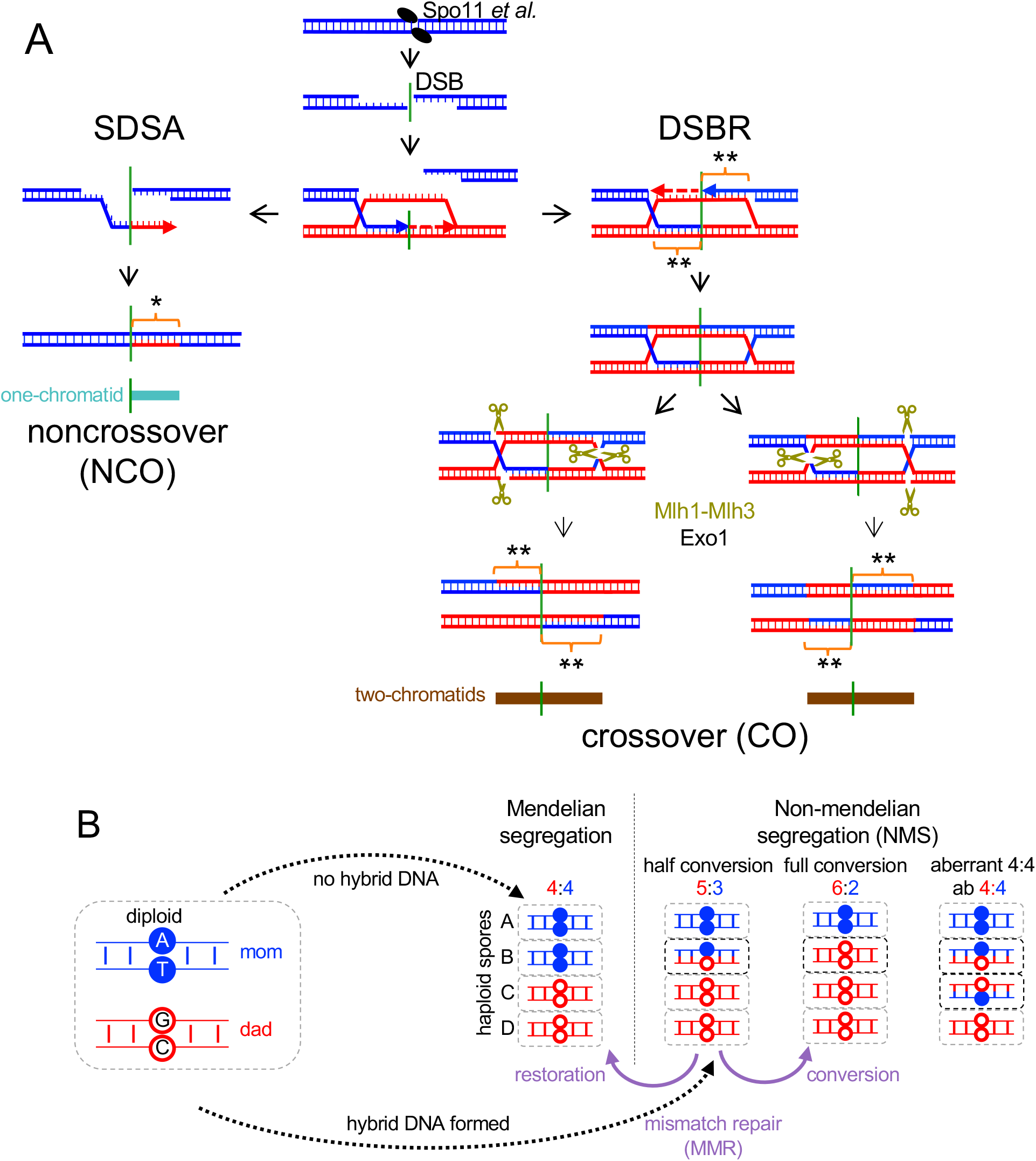
Current “dual-mechanism” model and marker segregation in tetrads. (A) Resection creates a 3’ ssDNA end that invades a homologous duplex and is extended by synthesis (arrow). SDSA – the invasion/extension intermediate is displaced and forms a NCO. DSBR – the second DSB end is captured and extended to form a double Holliday junction (dHJ) that resolves to form COs. Hybrid DNA in NCOs is continuous, on one chromatid and one side of the DSB (turquoise) and contains one “old” and one newly synthesized strand (*). COs contain hybrid DNA in two tracts (brown), one on each chromatid in opposite directions from the DSB, that contain two “old” parental strands (**). (B) If no hybrid DNA forms, markers segregate in a 4:4 ratio. Asymmetric hybrid DNA results in a 5:3 ratio (half conversion). Correction by MMR produces either 6:2 (full conversion) or 4:4 (restoration). Symmetric hybrid DNA produces an aberrant 4:4 ratio.

Dual-mechanism model intermediates contain DNA where the two strands are from different parents (Figure 1A), here called hybrid DNA. If homologs harbor different alleles, the resulting hybrid DNA will contain mismatches, and here will be called heteroduplex. The mismatches present in heteroduplex are detected genetically as departures from Mendelian segregation (4:4, based on the eight DNA strands present) in haploid meiotic products (Figure 1B). Heteroduplex on only one homolog (asymmetric heteroduplex) produces 5:3 or 3:5 marker segregation (half conversion) if left unrepaired, while heteroduplex on both homologs (symmetric heteroduplex) produces aberrant 4:4 (ab4:4) marker segregation. Mismatches in heteroduplex are recognized by mismatch recognition complexes (Msh2-Msh6 and Msh2-Msh3) and are corrected by the removal and resynthesis of strands over large regions, producing full conversions (6:2 or 2:6) or restoration to parental 4:4 ratios (Iyer et al., 2006; Spies and Fishel, 2015). Mismatch recognition also can trigger heteroduplex rejection, where helicases disassemble recombination intermediates that contain heteroduplex (Spies and Fishel, 2015). Deleting *MSH2* inactivates both heteroduplex rejection and MMR and thus preserves heteroduplex patterns, allowing inference of parental strand contributions from genotypes of spores in tetrads (Borts and Haber, 1987; Hunter and Borts, 1997; Martini et al., 2011).

The dual-mechanism model makes specific predictions regarding parental strand contributions to recombinants. In particular, NCOs formed by SDSA should be one-sided, with a continuous hybrid DNA tract that starts at the DSB site, while COs formed by DSBR should be two-sided, with two continuous hybrid DNA tracts that start at the DSB and extent in opposite directions. These predictions are challenged by the following observations:

- Frequent two-sided NCOs, with heteroduplex on both sides of the DSB on a single chromatid (Hoffmann and Borts, 2005; Jessop et al., 2005; Marsolier-Kergoat et al., 2018; Martini et al., 2011; McMahill et al., 2007; Porter et al., 1993).
- Mosaic heteroduplex, where heteroduplex tracts alternate with parental duplex (Crown et al., 2014; Marsolier-Kergoat et al., 2018; Martini et al., 2011). This can be produced by switching between chromatids as repair templates (McVey et al., 2004; Smith et al., 2007; Yeadon et al., 2001).
- Symmetrical hybrid DNA, detected as aberrant 4:4 (ab4:4) marker segregation (Figure 1B, Figure S1C). This can be formed by branch migration, where Holliday junctions move by exchanging base pairs, creating hybrid DNA on both recombining chromatids (Hamza et al., 1981; Holliday, 1964).
- Full conversion in mismatch correction-deficient cells (Williamson et al., 1985) or with markers that form poorly-recognized mismatches (Getz et al., 2008; Lichten et al., 1990; Nag et al., 1989).

Many of these observations were made in early studies that lacked the high density of heterozygous markers and knowledge of initiating DSB locations necessary for detailed mechanistic understanding. Recent studies, using *msh2*Δ mutants to preserve heteroduplex in hybrid budding yeast diploids with marker densities of about one per 140-170 nucleotides, provide compelling evidence for two-sided NCOs, template switching, branch migration and MMR-independent full conversion (Cooper et al., 2018; Marsolier-Kergoat et al., 2018; Martini et al., 2011; Oke et al., 2014). However, it remains uncertain the extent to which these noncanonical events can be generalized, in part because full analysis was precluded by uneven marker distributions and by the broad width and close spacing of many DSB hotspots (Pan et al., 2011).

To address this issue, we created a recombination reporter interval with uniformly spaced polymorphic markers and a single, narrow DSB hotspot, to map the origin, extent and structure of recombinants with unprecedented resolution. We find that many of the noncanonical events described above occur at high frequencies during meiotic recombination. Based on these findings, we suggest a novel mechanism for NCO and CO formation and differentiation.

## RESULTS

To study meiotic recombination at high resolution, we modified a recombination reporter interval that contains the *URA3* and *ARG4* genes and a single DSB hotspot (*his4::URA3-ARG4*; Figure 2A; Jessop et al., 2005). The modified locus (*his4::URA3-ARG4-SNPs*; Figure 2A, File S1.2) contains sequence changes every 96 ± 29 bp (mean ± S.D) for 1 kb to the left of the DSB hotspot and 2 kb to the right, with a lower polymorphism density for another 1.5 kb to the left and 1.1 kb to the right. The changes preserve wild-type coding and create flanking *Xmn*I site polymorphisms for molecular CO scoring. Flanking drug resistance markers allow genetic CO scoring.

**Figure 2.**
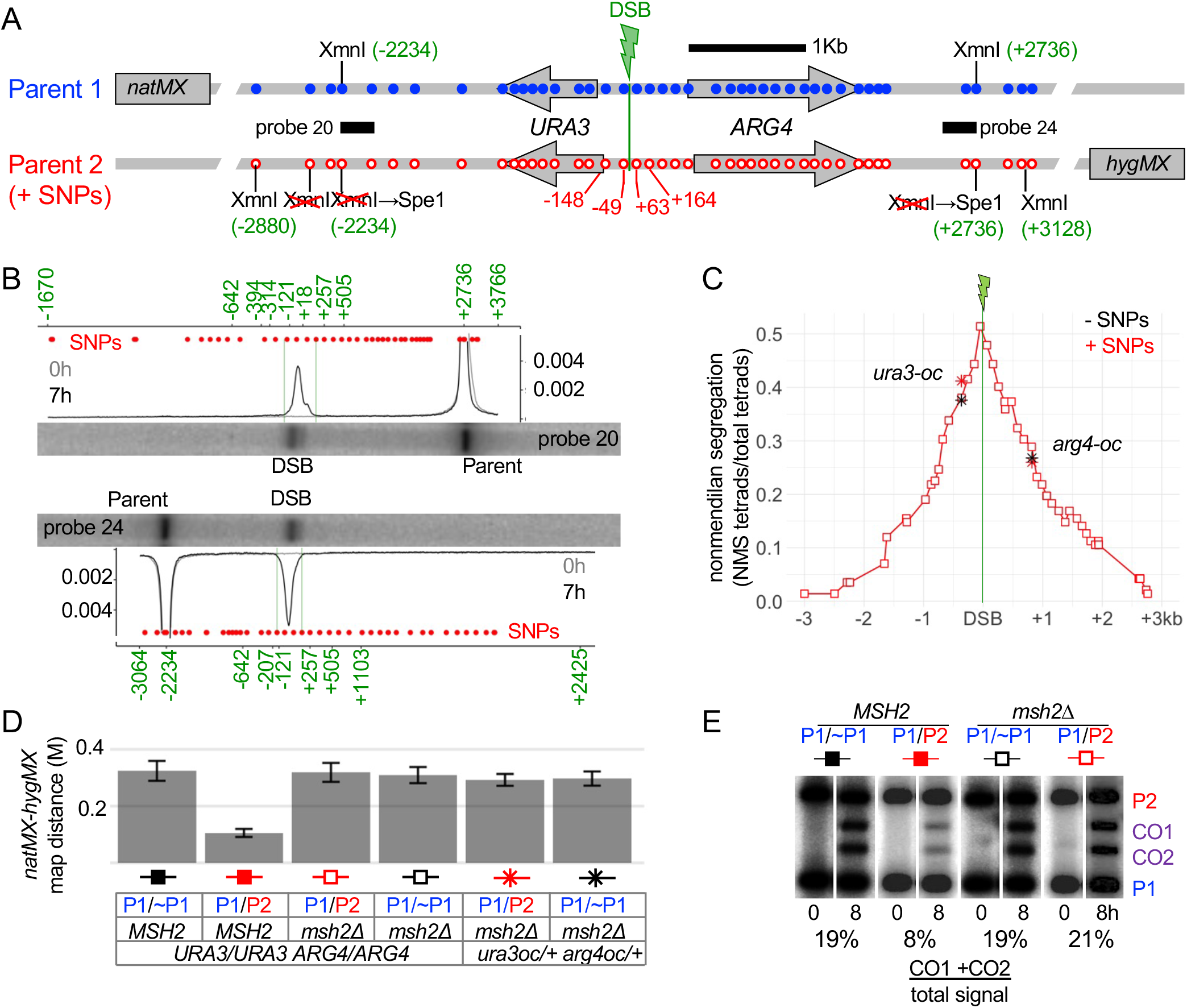
Characteristics of the interval studied. (A) The URA3-ARG4 recombination interval, showing polymorphic markers (blue—wild type; red– polymorphisms). Marker locations are relative to the DSB centroid (File S1.2). (B) DSBs are tightly focused. Southern blots of XmnI/SpeI digests of meiotic DNA from a resection-deficient rad50S strain (see A for probes). Plots are signal/total lane signal. Red dots—marker locations; green numbers—size standards (Figure S2; high resolution gels in Figure S2C, D). (C) Nonmendelian segregation (NMS). Red squares—NMS determined by sequencing. Black stars—NMS for ura3-oc (−364) and arg4-oc (+819) in strains lacking polymorphisms from -2234 to +2736. Red stars—NMS for the same mutants in strains heterozygous for the full polymorphism set. (D) Heterozygosity reduces COs in the natMX:hygMX interval in wild type but not in msh2Δ. Map distances (Morgans) from tetrad analyses; P1 and P2 are as in panel A; ∼P1 lacks sequence polymorphisms from -2234 to +2736 (Figure S2A). Error bars—standard error. (E) Heterozygosity reduces COs in wild type but not in msh2Δ. XmnI digests of DNA from cells before (0h) and after meiosis (8h), probed with ARG4 sequences.

Meiotic DSBs form in this interval in 17.5% of chromatids (Figure 2B) and are tightly focused, with 84% of DSBs in a 112 nt region between the two central markers (−49 nt and +63 nt relative to the DSB centroid). All DSBs detected in the interval were located in a 312 nt region flanked by the next two markers (−148 to +164, Figure 2B, Figure S2B).

Marker segregation in 142 tetrads from a MMR-deficient *msh2*Δ diploid was scored by high-throughput sequencing of amplicons from spore colony DNA (File S1.3, File S2). Non-mendelian segregation (NMS) occurred in 77% of tetrads, consistent with DSB levels; 3/4 of NMS were half conversions (File S1.4). Markers immediately adjacent to the DSB centroid showed maximal NMS (51% and 48%, Figure 2C, File S1.4), which decayed exponentially with distance (median NMS ∼780 nt; Figure S2E). In the absence of MMR, the high polymorphism density did not affect recombination outcomes: NMS for *ura3* and *arg4* ochre mutations and COs in the interval were similar in *msh2*Δ strains with a fully heterozygous central or a homozygous interval (Fig. 2C-E, File S1.5 and S1.6). In MMR-competent *MSH2* strains, heterozygous polymorphisms reduced COs ∼3-fold (Figure 2D, E). This substantial effect, for a highly heterozygous interval in an otherwise homozygous genome, stands in contrast to the lower genome-wide CO reductions (1.3-1.4 fold) seen in hybrid strains with genome-wide heterozygosity (Cooper et al., 2018; Marsolier-Kergoat et al., 2018; Martini et al., 2011). It is likely that in *MSH2* strains with genome-wide heterozygosity, MMR decreases COs in highly heterozygous regions, and CO homeostasis increases COs in less heterozygous regions (Cooper et al., 2018; Martini et al., 2006; Thacker et al., 2014).

38% (55/142) of tetrads contained a CO involving two chromatids, and 23% (32/142) contained a single NCO, most of which (28/32) involved conversion of one chromatid. 15% of tetrads (22/142) contained events involving three or four chromatids. About half of these (10/22) must have involved DSBs on more than one chromatid, while the remainder (12/22) could have been produced by more than one DSB or by resolution of a multichromatid JM initiated by a single break (Figure S1A, File S1.1). Because of this uncertainty, we focused on the ∼3/4 of recombinant tetrads that appeared to come from a single initiation event.

### Similar patterns of end invasion and extension among noncrossovers and crossovers

The dual-mechanism model predicts that NCOs contain hybrid DNA on one side of the initiating DSB, while COs will contain hybrid DNA formation on both sides (Figure 1A). The heteroduplex patterns we observed do not support this prediction (Figure 3). Instead, NCOs and COs have remarkably similar patterns of heteroduplex. Two-thirds of NCOs (22/32) have NMS tracts on both sides of the DSB, and about one-third of COs (20/55) have a single NMS tract on one side of the DSB (Figure 3D). This similarity between NCOs and COs extends to total NMS tract lengths (Figure 3E) and to half-tract lengths (distance from DSB to NMS tract end; Figures 3F; File S1.7).

**Figure 3.**
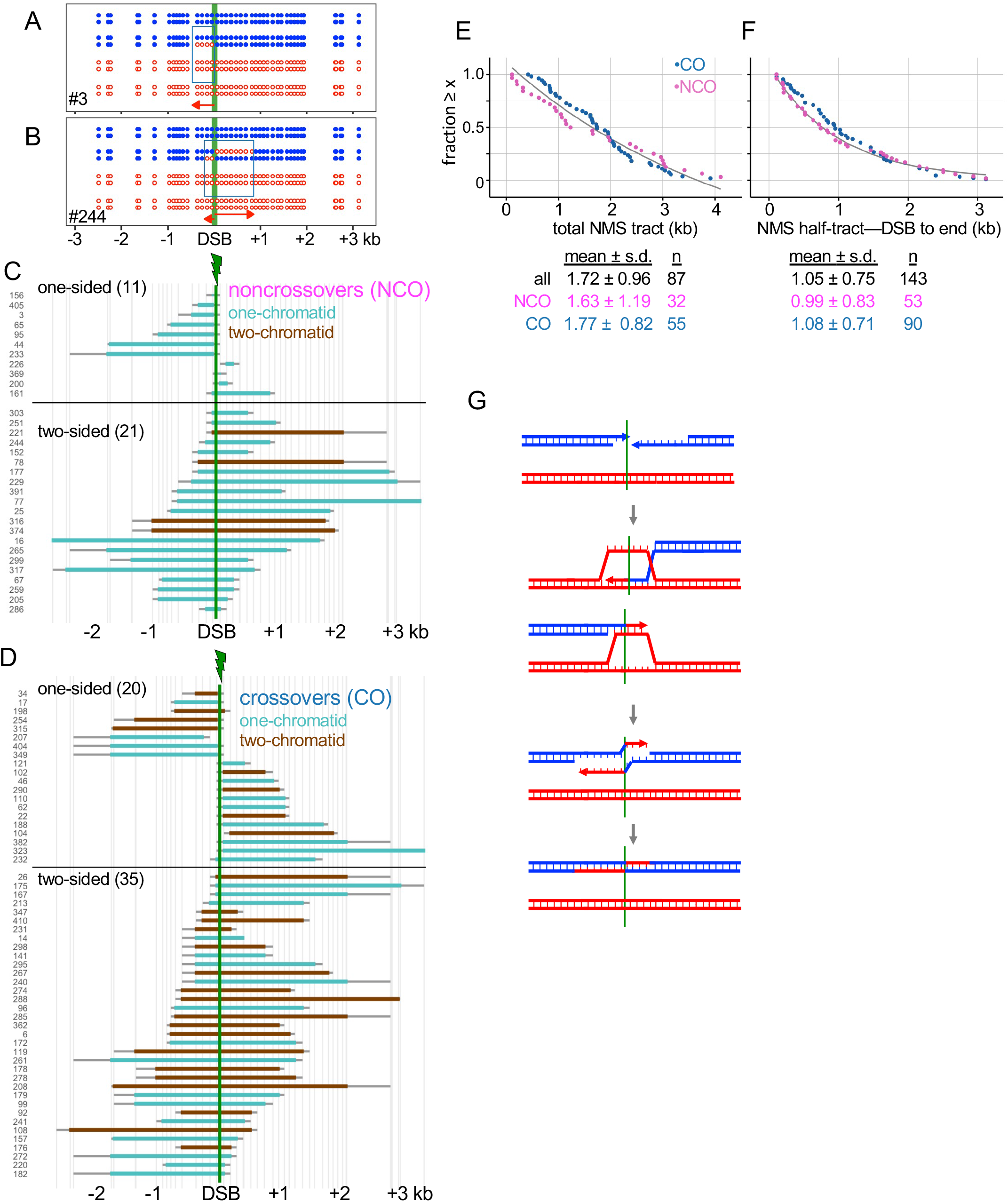
Similar patterns of end invasion and extension in NCOs and COs. (A) NCO tetrad with one-sided NMS. (B) NCO tetrad with two-sided NMS. (C) NMS tracts in NCOs. Turquoise—heteroduplex on one chromatid; brown—heteroduplex on two chromatids (see Figure 1A). Vertical axis–tetrad identifiers; vertical lines—marker positions. Thick colored bars and thin gray bars indicate minimum and maximum NMS tracts, respectively. (D) NMS tracts in COs; color code as in C. (E) Rank order plot of NMS tract lengths for crossovers (blue) and noncrossovers (purple), calculated using the average between minimum and maximum NMS tracts. Black line—exponential decay curve fit to all events. (F) NMS half-tract lengths, distance from the initiating DSB (midpoint between closest flanking markers) and the NMS tract end (midpoint between the last converted and the first unconverted marker); colors as in E. (G) Double SDSA. Both DSB ends invade a homolog, extend, are displaced, and anneal to form a two-sided NCO. Simultaneous invasion of two homologs is shown here, but invasion could occur sequentially and could involve a single chromatid.

Two-sided NCOs can be produced by a process called double SDSA, in which the two DSB ends both invade, extend and are displaced before annealing with each other (Figure 3G). Apparent one-sided COs might be two-sided events where one hybrid DNA tract is shorter than ∼100 nt, and thus might not have included a marker. If every one-sided CO contains a second “invisible” hybrid DNA tract, then 18% of CO NMS tracts are expected to be <100 nt. However, extrapolation of observed NMS half-tract lengths predicts that only ∼8% should be ≤ 100nt (Figure S3B). Thus, it is likely that many of COs scored as one-sided contain hybrid DNA on only one side of the DSB.

### Template switching is common among both crossovers and noncrossovers

It has been suggested that meiotic recombination involves multiple rounds of invasion, D-loop disassembly and re-invasion (De Muyt et al., 2012; Kaur et al., 2015). If switching between the homolog and sister chromatids occurs during this process, mosaic hybrid DNA will be produced (Figure 4A). We counted as mosaic only those NMS tracts that had at least one half-conversion (5:3 or 3:5) segment and at least one segment with two or more markers in a parental (4:4) configuration (Figure 4B). Single marker 4:4 segregations were not counted, as they can also be produced by *MSH2-*independent short-patch mismatch repair (Coic et al., 2000; Crown et al., 2014; Fleck et al., 1999). This suggestion is supported by the similar recovery, among NCOs, of 4:4 and 6:2 single-marker segregations inside heteroduplex tracts (File S1.12). Other marker segregation patterns (full conversion and symmetrical heteroduplex) that mostly occur in COs also were not counted; these will be discussed below.

**Figure 4.**
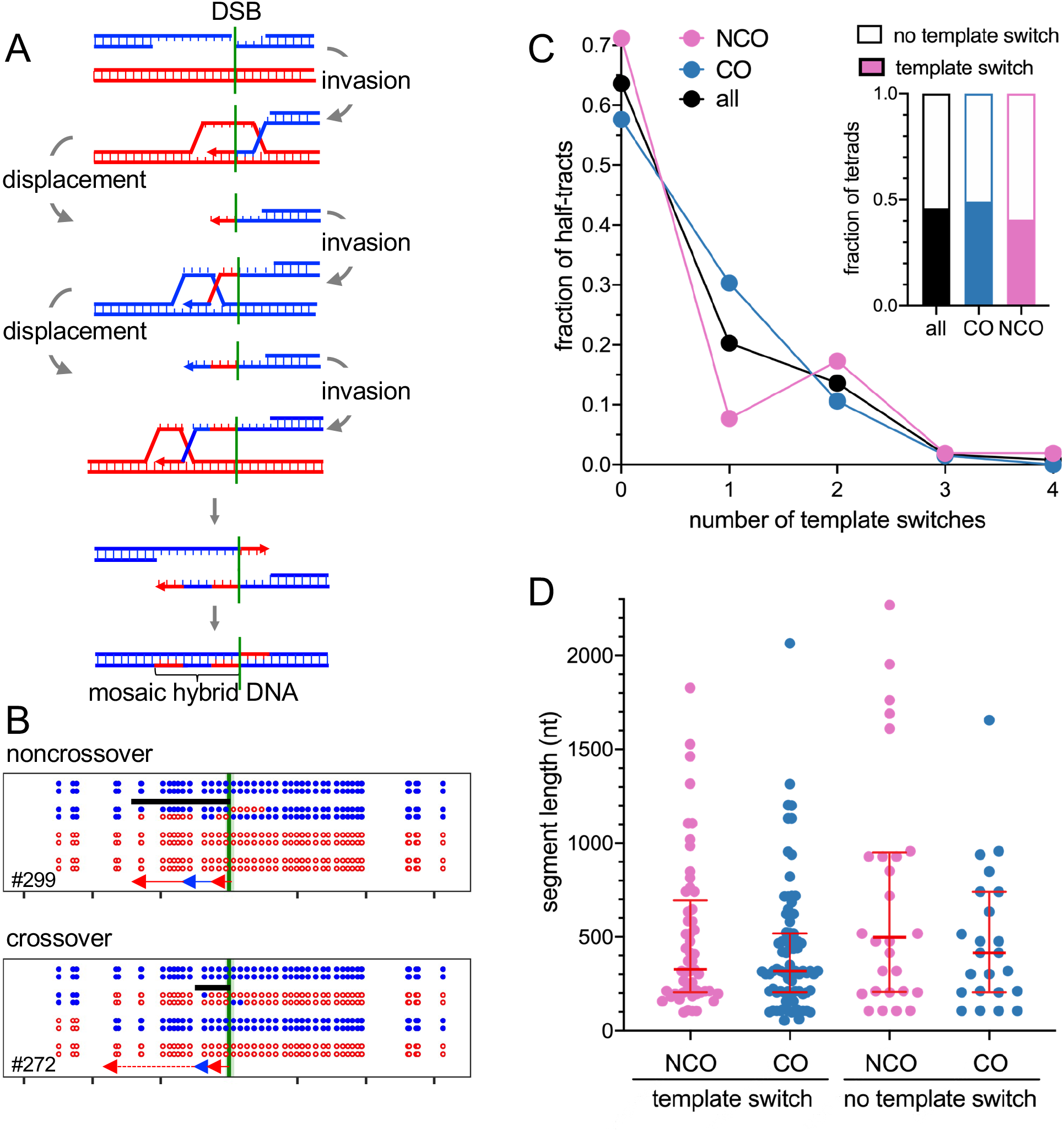
Template switching is common in both COs and NCOs. A) Multiple rounds of invasion, extension, and displacement can form mosaic hybrid DNA. (B) Examples of mosaic heteroduplex among NCOs and COs. Black bars–minimum extent of mosaic heteroduplex. The NCO first invaded the homolog, while the CO first invaded the sister chromatid. The second parental segment in the CO (dotted line) could have involved either invasion of the sister chromatid or branch migration (see Figure 5). (C) Frequent template switching in COs and NCOs. Inset—fraction of tetrads with template switching. Tetrads where interpretation was uncertain are counted as “no template switch”. Main graph—number of template switches per NMS half-tract. (D) Length of segments in NMS half-tracts with or without template switching. Red lines–median and quartiles. None of the distributions are significantly different from the others (p>0.05, Mann-Whitney test).

NCOs and COs displayed similar levels of mosaic heteroduplex (∼40-50%; Figure 4C, File S1.4) and similar frequencies of template-switching (mean of 0.6 ± 0.1 switches/tract ± S.E.M. for both; Figure 4C) and switch segment lengths (median of 327 versus 319 nt/segment; Figure 4D). Non-mosaic NMS tracts also were similar lengths in NCOs and COs (Figure 4D).

The DSB-adjacent segment in mosaic heteroduplex indicates which chromatid was first invaded (4:4, sister chromatid; 5:3, homolog; Figure 4B). About 1/3 of DSB-adjacent segments were parental (20/64 in NCOs, 45/110 in COs, File S1.4). This indicates that DSB ends invade the one sister and two homolog chromatids with equal likelihoods, as has been previously suggested (Goldfarb and Lichten, 2010; Oh et al., 2007).

The presence of mosaic heteroduplex in roughly half of both COs and NCOs indicates that template switching is frequent during meiotic recombination. This is a minimum estimate, as genetically invisible template switches—for example, from one homolog chromatid to the other, or to the sister chromatid as a final switch—would not have been detected.

In summary, both DSB ends invade and extend in a majority of both NCOs and COs, and a similar fraction of NCOs and COs undergo template switching (Figure S4), consistent with the suggestion that similar mechanisms form hybrid DNA in both NCOs and COs.

### Branch migration is frequent among crossovers

dHJ resolution produces a transition from hybrid DNA to parental sequences. A central feature of the DSBR model is that all hybrid DNA should be between the two resolution points (Figure 1A). Consequently, hybrid DNA should be on one side of the DSB on one chromatid, to other side of the DSB on the other (Figure 1A).

Remarkably, the vast majority of COs did not conform to these predictions (Figure 5, File S1.10). In half of COs (28/55), both exchange points were on the same side of the DSB (Figure 5A, D), and heteroduplex tracts on a single chromosome often spanned the DSB (example in Figure 5C). In addition, many CO exchange points were separated from DSB-adjacent heteroduplex by two or more markers in a parental configuration (Figure 5A, Figure S5A). In some cases (7/28), the two exchange points flanked a stretch of symmetrical heteroduplex (hybrid DNA present on both chromatids; aberrant 4:4 segregation; Figure 5A, File S1.10). Detection of such heteroduplex was limited by larger inter-marker intervals distant from the DSB (Figure 2A), and by the destruction of CO-adjacent heteroduplex by strand processing, which will be discussed in the following section.

**Figure 5.**
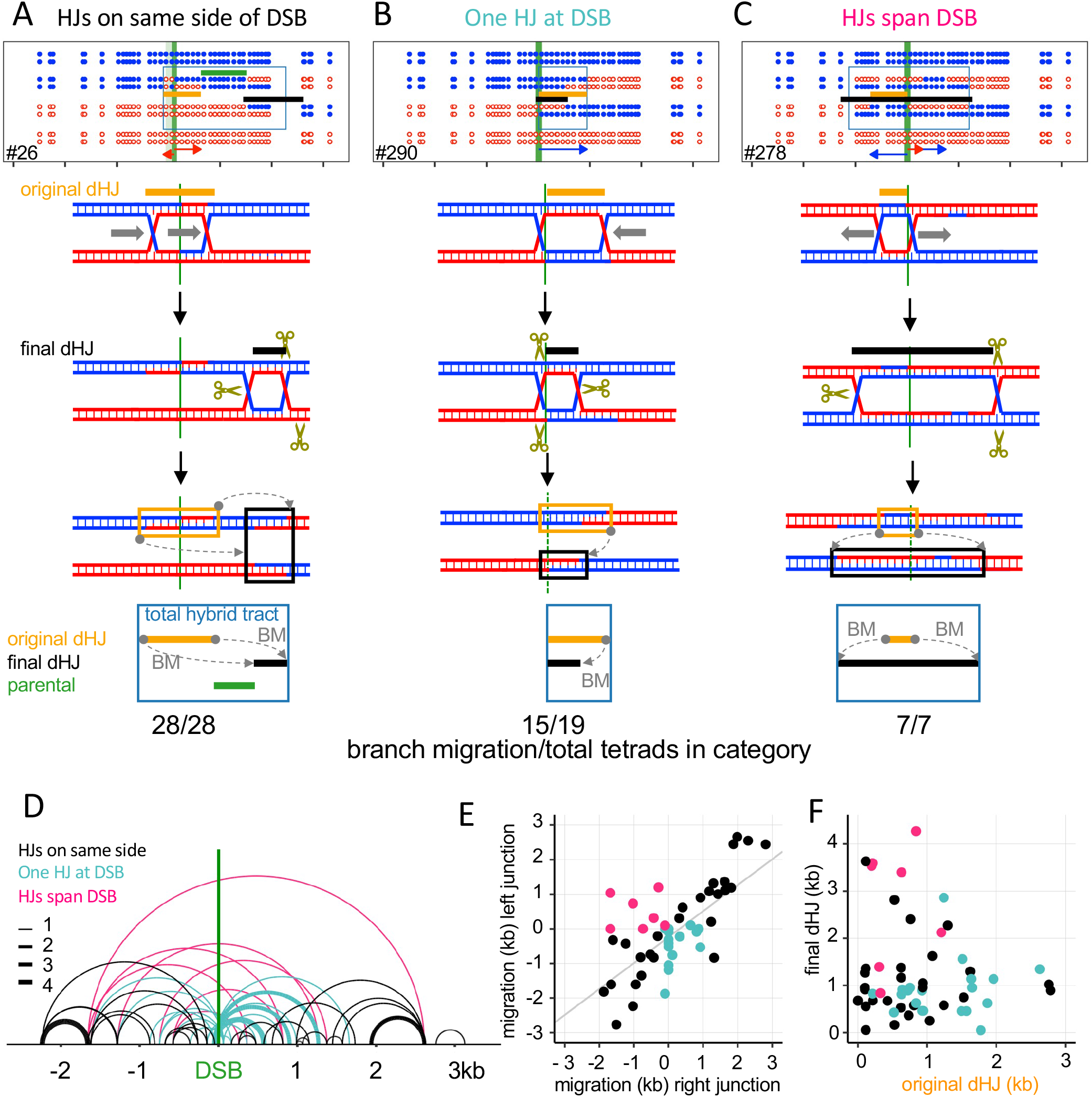
Crossovers frequently involve Holliday junction branch migration. Top—COs classified with regards to final HJ locations (crossover points). Middle – an example for each category. Bottom – proposed branch migration mechanism producing the final marker pattern. Tan– inferred original HJs; black –final HJs; green –parental sequences between original and final dHJs; blue boxes— total hybrid DNA. Grey arrows (BM) – direction and extent of branch migration. The fraction of each category displaying branch migration is given below. (A) Both final HJs are on the same side of the DSB interval. (B) One HJ is in the DSB interval, and the other to one side. (C) HJs are on opposite sides of the DSB interval. In this example, both junctions have moved outward. (D) Final dHJ locations. Arcs connect the two HJs. Colors correspond to categories in A-C; line thickness indicates the number of events. (E) Distance and direction of migration by the two HJs. Negative and positive values denote left- and rightward migration; colors as in D. (F) distances between original and final dHJ midpoints; colors as in D.

These noncanonical heteroduplex patterns can be produced by branch migration (Holliday, 1964). Movement of a HJ through parental duplex creates symmetrical hybrid DNA (ab4:4). Codirectional movement of the second HJ in a dHJ restores this to the original 4:4 configuration, while retaining a patch of symmetrical hybrid DNA between the two junctions (Figure S1Ci, Figure 5A). Migration of a single junction through a region of hybrid DNA transfers hybrid DNA from one chromatid to the other (Figure S1Cii). This can result in a CO where hybrid DNA switches from one homolog to the other (Figure S1Ciia). These patterns are prominent among the COs where both Holliday junctions are on one side of the DSB (Figure 5A; File S1.10).

Signals of branch migration are also present in COs where the final HJs were on opposite sides of the DSB (Figure 5C, D; 7/7), and in COs where one final HJ was inside the DSB region (Figure 5B, D; 15/19). In total, the vast majority (87%) of COs displayed hallmarks of branch migration. In contrast, a much smaller fraction of NCOs (3/22, File S1.10) showed evidence of branch migration, consistent with most NCOs not involving dHJ formation and resolution.

In many cases, initial junction positions could be inferred from heteroduplex patterns by assuming that the initial intermediate was a dHJ, and thus could calculate the direction and distance each junction migrated (Figure 5E, Figure S5A-C, File S1.10). In some cases only one junction moved; in others the two junctions moved in opposite directions, or both moved in the same direction but for different distances (Figure 5E). Initial and final inter-junction distances often differed from each other (Figure 5E, Figure S5D).

Intermediates with two cross-strand junctions, including dHJs, are topologically constrained by the helical turns between the two junctions (Kaur et al., 2015), and movement of one junction will drive similar movement of the other, thus preserving the original inter-junction distance. In most COs, the two junctions appear to have moved different distances and/or in different directions (Figure 5E). This would require relief of topological constraints, either by topoisomerase activity or by junction-nicking during resolution. Alternatively, it is possible that much of the heteroduplex in COs forms by topologically unconstrained mechanisms. This will be discussed further below.

### Crossovers display resolution-associated strand processing

Crossovers frequently contained full conversions (6:2 segregation), even though standard MMR is absent from the *msh2*Δ strains used. As with template switching, we only considered full conversion of two or more adjacent markers, given that single-marker full conversion can be produced by *MSH2*-independent short-patch repair (Coic et al., 2000; Crown et al., 2014; Fleck et al., 1999). Full conversion was prominent among crossovers (19/55 crossovers, Figure 6A, Figure S6A, File S1.11), rare among noncrossovers (1/32; Figure S6A), and was associated with potential HJ resolution sites (Figure 6A, File S1.11, File S2). This might be expected if HJ resolution-associated nicks were initiation points for removal and resynthesis of one heteroduplex strand (Figure 6; Kulkarni et al., 2020; Marsolier-Kergoat et al., 2018). Another six additional tetrads had full conversion of a single marker at a resolution point, and thus are potential candidates for resolution-associated conversion (Figure S6A). Full conversion can be produced by repair of a dsDNA gap (Johnson et al., 2019; Prieler et al., 2021; Szostak et al., 1983; Figure S1D), but most of the tracts had at least one end outside of the DSB region, and tracts frequently were separated from the DSB region by markers in a parental configuration (File S1.11, Supplementary File 2). It is therefore unlikely that most are products of double strand gap repair (see Discussion). Another 7 COs contained other indications of resolution-associated strand processing (Figure 6B, Figure S6A, File S1.11,1), including tetrads where symmetrical heteroduplex was converted to asymmetrical heteroduplex. Processing tracts were short (median ∼507 nt for multiple marker events; Figure 6C), so it is possible that some strand processing events were not detected.

**Figure 6.**
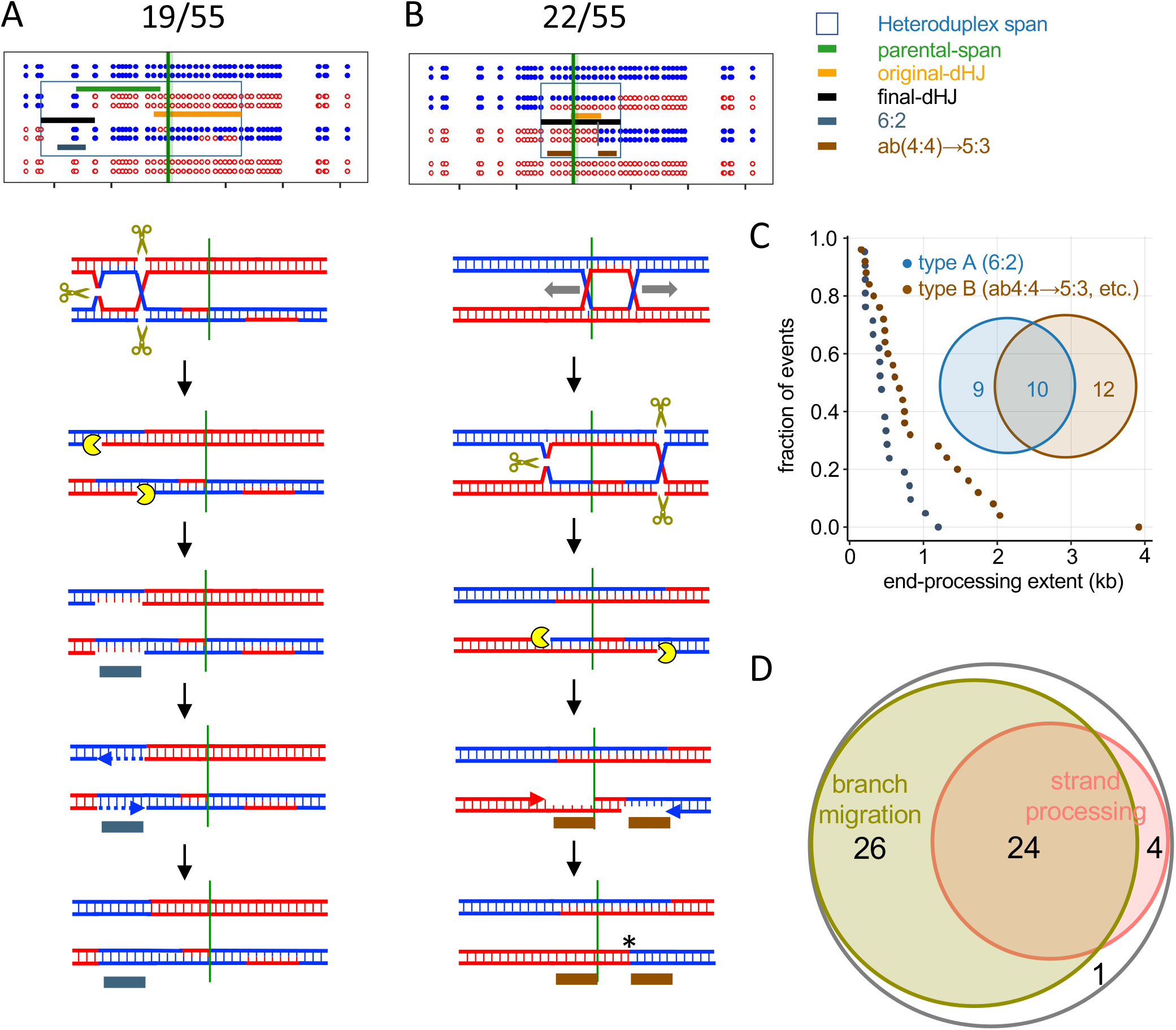
Crossovers frequently involve resolution-associated strand processing. (A) CO where end-processing produces 6:2 segregation. dHJ resolution produces nicks that are converted to ssDNA gaps; repair synthesis results in full conversion (Stahl and Foss, 2010). (B) CO where end-processing converts ab(4:4) to 5:3 segregation, resulting in three apparent CO points. Colored indicators as in Figure 5, with the following additions: grey—6:2 tract produced by end processing; brown—5:3 tract produced by end processing. The fraction of COs in each category is listed. (C) Rank order plots of strand processing tract lengths in types A and B. Only tracts of 2 or more markers were tabulated; curves including single-marker events are in Figure S6B. Median tract lengths: type A— 412 nt; type B—679 nt. Inset Venn diagram--tetrads with type A and type B strand processing. (D) Number of CO tetrads displaying branch migration, end processing, or both. One tetrad was too complicated to be scored.

Taken together, about half of COs display signals of resolution-associated strand processing, and the vast majority of these also displayed hallmarks of branch migration (Figure 6D). The frequent overlap between template switching, branch migration and resolution associated strand processing often produced COs with highly complex parental marker segregation patterns (File S2). In all, 54/55 COs displayed evidence for one or more of these processes, a remarkable departure from the predictions of the DSBR model.

## DISCUSSION

Since the dual-mechanism model was first proposed (Allers and Lichten, 2001a), most studies of meiotic recombination have been interpreted in its context. Recent reports documenting incompatible heteroduplex patterns challenge this model (Marsolier-Kergoat et al., 2018; Martini et al., 2011; Peterson et al., 2020), but leave open the question of the degree to which such “non-standard” events are typical of meiotic recombination. The recombination reporter we developed, with close, regularly spaced markers and a single tightly focused DSB hotspot, has allowed characterization of individual events with unparalleled precision, to show that these “non-standard” processes are the rule. Below, we discuss the implications of these findings, including a model for meiotic recombination that accounts for features shared and different between NCOs and COs.

### Similar end invasion, extension and mosaic heteroduplex patterns in NCOs and COs suggest a common mechanism

We find that NCOs and COs display similar patterns of heteroduplex, both with regards to one-versus two-sided events (Figure 3) and mosaic heteroduplex (Figure 4). In particular, a substantial majority (70-80%) involved homolog invasion by both DSB ends (two-sided events), multiple invasions by a single DSB end (template switching), or both (Figure S4). Thus, multiple rounds of repair template invasion and disengagement are the rule, rather than the exception, during all types of meiotic recombination.

We first discuss these findings in the context of NCOs. Simple SDSA produces a single, one-sided hybrid DNA tract, but two thirds of NCOs contain heteroduplex on both sides of the DSB, most (17/21) with heteroduplex on a single chromatid and a switch in heteroduplex phase at the DSB (Figure 3B, File S2). It is likely that these are produced by double SDSA (Figure 3H). Thus, at least 2/3 of NCOs involve invasion by both DSB ends, and this fraction is likely greater, as two-ended invasions where one end invaded a sister chromatid would have been scored as one-sided.

A substantial fraction of NCOs (∼40%) contained mosaic heteroduplex (Figure 4C, File S1.10). While mosaic heteroduplex could be produced by Msh2-independent repair, the absence of full conversion segments inside NMS tracts (File S1 Sheets 11, 13, 14) indicates that this is not likely. We therefore conclude that mosaic heteroduplex is produced by template switching (Figure 4A). As was noted above, many template switching events might not have been detected om our analysis, making it likely that the majority of NCOs undergo multiple rounds of invasion, disassembly, and re-invasion.

Other studies report that two-sided NCOs are a minority class (Gilbertson and Stahl, 1996; Hoffmann and Borts, 2005; Jessop et al., 2005; Merker et al., 2003; Porter et al., 1993), and report mosaic heteroduplex frequencies lower than those we observe (Marsolier-Kergoat et al., 2018; Martini et al., 2011). This disparity may be due to uncertainty regarding DSB locations and/or limits on the ability to detect hybrid DNA, due to low or non-uniform marker densities (c.f. Marsolier-Kergoat et al., 2018; Figure S3A).

Remarkably, COs displayed heteroduplex patterns similar to those in NCOs. A similar fraction of COs and NCOs contained heteroduplex on only one side of the initiating DSB (Figure 3C,D), and about half of COs with two-sided heteroduplex contain all heteroduplex on a single chromatid (Figure 3D). Neither outcome is predicted by the DSBR model. In addition, COs and NCOs had similar heteroduplex tract length distributions (Figure 3E, F), a finding not expected in the dual mechanism model, in light of its different mechanisms for hybrid DNA formation.

COs also contained mosaic heteroduplex at levels and with patterns comparable to those in NCOs (Figure 4C, D). Since mosaic hybrid DNA is formed by multiple rounds of invasion and end-primed synthesis (Figure 4A), the mosaic heteroduplex found in COs must be formed by annealing between an “old” and a “new” strand. This runs counter to the DSBR model, where all hybrid DNA in dHJs, and thus in COs, contains two “old” DNA strands (Figure 1A). For the same reason, we believe it unlikely that NCOs, at least those that contain mosaic heteroduplex, are be formed by dHJ dissolution.

Our finding that 1/3 of mosaic heteroduplex involves an initial invasion of the sister (File S1.9) challenges the view, largely based on characterization of dHJs in budding yeast (Schwacha and Kleckner, 1994), that inter-sister recombination is disfavored during meiosis. While it is clear that meiotic recombination does not have the strong preference for inter-sister events seen in mitotic cells (Bzymek et al., 2010; Kadyk and Hartwell, 1992; Lichten and Haber, 1989), our data support previous conclusions that DSB ends have similar likelihoods, on a per-chromatid basis, of engaging sister and homolog chromatids (Goldfarb and Lichten, 2010). Frequent engagement of the sister chromatid during meiosis has also been proposed for mouse and fission yeast (Cole et al., 2014; Cromie et al., 2006), and thus may be a feature common to meiotic recombination in all organisms. The observed bias towards interhomolog dHJs may instead reflect the preferential disassembly of intersister recombination intermediates (De Muyt et al., 2012; Goldfarb and Lichten, 2010; Kaur et al., 2015; Oh et al., 2007; Sandhu et al., 2020; Tang et al., 2015).

In summary, the high frequency of mosaic heteroduplex in both COs and NCOs, combined with other features of heteroduplex common to NCOs and COs, suggests that most hybrid DNA in NCOs and COs is formed by a common mechanism that need not involve dHJ formation.

### Branch migration is intrinsic to crossover formation

While the above-discussed processes are common to all meiotic recombination, others appear to be CO-specific. The most prominent of these, junction movement, occurs in the vast majority (85-90%) of COs but only a small fraction of NCOs (Figure 5; File S1.1 and S1.10). About half of COs (28/55) contained both final junctions on the same side of the DSB (Figure 5A, D). These two junctions often were separated from the DSB by a region of 4:4 marker segregation, and they often flanked symmetrical heteroduplex (Figure 5, Figure S5A). These configurations can be produced by branch migration of both junctions in the same direction (Allers and Lichten, 2001b; Marsolier-Kergoat et al., 2018; Figure S1D), but are incompatible with other mechanisms, such as D-loop or bubble migration (Ferguson and Holloman, 1996; Hoffmann and Borts, 2005), that involve continuous DNA synthesis.

The near-ubiquity of branch migration suggests that it is an essential step in CO formation. Even the canonical DSBR model requires branch migration to facilitate closure of the nicks present at cross-strand junctions (Figure S7A; Szostak et al., 1983). However, the extent of junction movement we observe (median of at least 0.66 kb; Figure S5D) is considerably greater than that required for ligation, indicating that it serves other functions.

To account for these phenomena, we suggest that during both NCO and CO formation, early intermediates are engaged by helicases at or soon after strand invasion (Allers and Lichten, 2001b; De Muyt et al., 2012; Lao et al., 2008; Figure 7). An intermediate that is not CO-designated undergoes helicase-mediated disassembly, followed by either additional rounds of invasion and extension, or by annealing with the other DSB end to form a NCO (De Muyt et al., 2012). CO-designation of an intermediate occurs when branched structures are stabilized by ZMM proteins, most likely the Zip2-Zip4-Spo16 heterotrimer (De Muyt et al., 2018), and helicase activity is redirected towards branch migration, which confers further stability by converting D-loops into structures with two crossed-strand junctions without a need for second end capture or ligation (Figure 7B, Lao et al., 2008; Oh et al., 2007). Branch migration also re-exposes ssDNA ends, like those liberated by D-loop disassembly in SDSA, that can participate in further rounds of strand invasion and extension, before they anneal with the other DSB end and create a four-armed intermediate (Figure 7, Figure S7B-D). Thus, CO formation involves two mechanistically distinct and potentially uncoupled processes, one that forms dHJs, and the other that forms DSB-associated heteroduplex (Figure 7B, Figure S7). We refer to this model, first proposed by Lao et al. (Lao et al., 2008) with features from previous studies (Allers and Lichten, 2001b), as the disassembly/migration-annealing (D/M-A) model.

**Figure 7.**
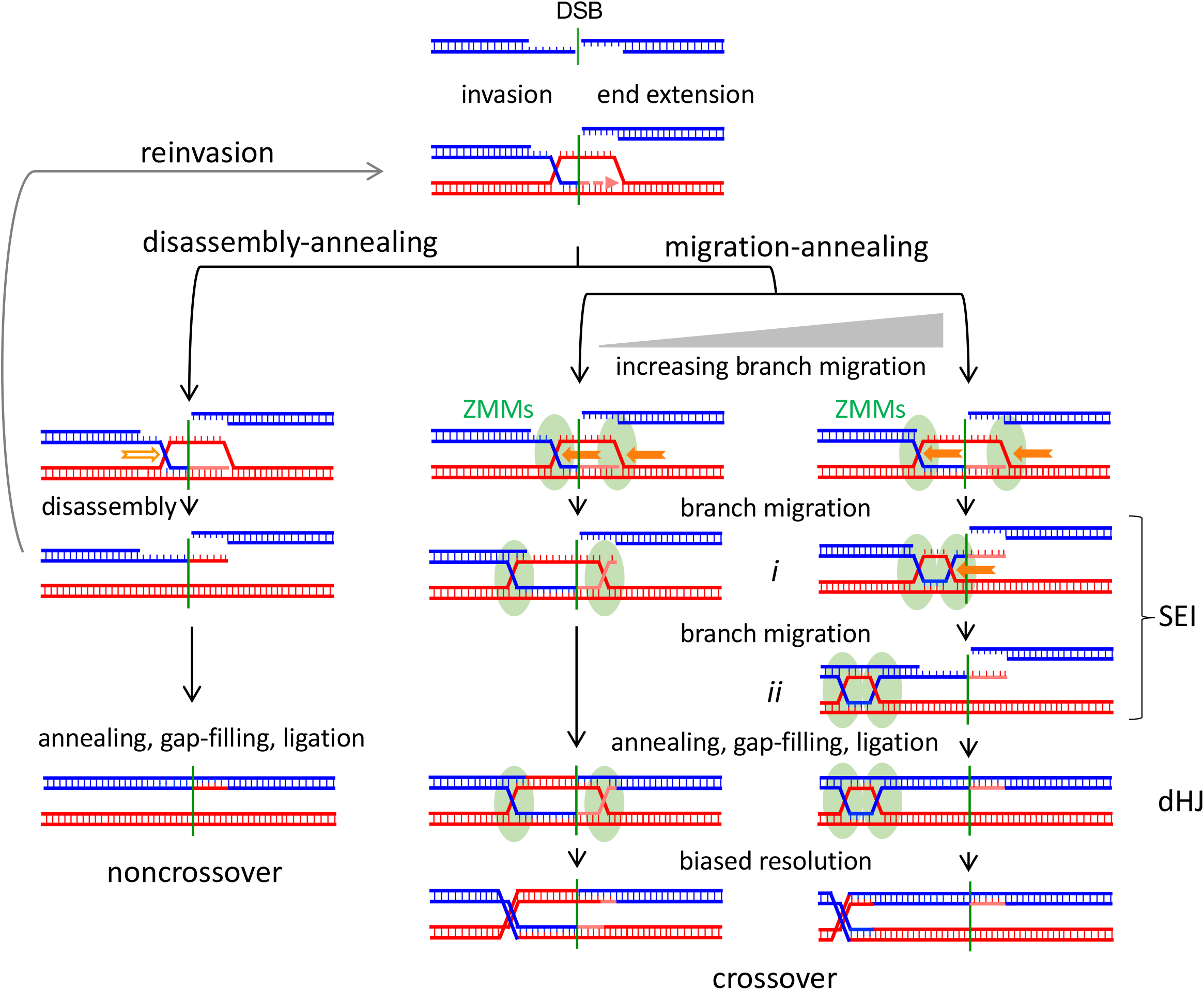
Disassembly/migration-annealing model for meiotic recombination. This model incorporates features of previous proposals (Allers and Lichten, 2001a, b; Lao et al., 2008). Left–helicase-mediated branch migration (hollow arrow) disassembles a D-loop, followed either by reinvasion or by end annealing to form a NCO. Center, right–CO formation. D-loop branches are captured by ZMM proteins and undergo helicase-mediated branch migration (solid arrows). Migration for different distances creates a three-arm single-end intermediate (SEI; Hunter and Kleckner, 2001) with a topologically closed bubble (i) or a dHJ (ii). Migration is drawn as unidirectional but may well be reversible. Annealing with the other DSB end and further processing produces a four-armed dHJ that is resolved as a CO. End reinvasion before annealing is also possible but is not drawn here. Center– limited migration can produce a DSBR-like intermediate. Migration after annealing is also possible (Figure S7A), as are different hybrid DNA configurations relative to the CO and DSB (Figure S7B).

While current data are difficult to reconcile with the dual mechanism model, the D/M-A model accommodates the following:

- Because both NCOs and COs involve annealing between an “old” strand exposed by resection and a “new” strand formed by end-primed synthesis, NCOs and COs should display similar patterns of heteroduplex (c.f. Figures 3, 4), and mutants that affect heteroduplex tract lengths should affect NCOs and COs similarly (c.f. Duroc et al., 2017; Vernekar et al., 2021).
- The three-arm intermediates formed by branch migration (Figure 7i and ii) corresponds to an observed structure, called an SEI (single-end intermediate; (Hunter and Kleckner, 2001). SEIs are as ZMM-dependent as are four-arm dHJs (Börner et al., 2004), and the two intermediates accumulate to similar extents in resolution-defective *ndt80* mutants (Shinohara et al., 2008).
- The suggestion that NCO formation and the SEI to dHJ transition occur by annealing (Figure 7) is consistent with the strong reduction in NCOs and four-arm dHJs, but not in SEIs, seen strand annealing-defective *rad52*Δ mutants (Lao et al., 2008).
- dHJ migration leaves sequences between initial and final dHJ positions in a parental configuration (Figure 7), consistent with previous findings that ∼1/4 to ∼1/3 of COs display parental marker segregation between the CO and the initiating DSB (Allers and Lichten, 2001b; Jessop et al., 2005).

Spontaneous dHJ migration is expected to be slow *in vivo*, especially in the presence of mismatches (Panyutin and Hsieh, 1993, 1994). The finding that branch migration can cover substantial distances (Figure 5) suggests that it is actively driven. There are two leading candidates for this function. The first is the conserved meiosis-specific Mer3/Hfm1 helicase, a ZMM protein that is required for normal dHJ-JM and CO formation (Börner et al., 2004; Mercier et al., 2005; Pezza et al., 2006; Wang et al., 2009). The second is the Bloom helicase (Sgs1 in budding yeast), part of a complex with Top3 and Rmi1. Sgs1 is best known for its anti-crossover activity (De Muyt et al., 2012; Ira et al., 2003) but also is required for ZMM-dependent meiotic COs (De Muyt et al., 2012; Zakharyevich et al., 2012). It is possible that Sgs1, perhaps in a post-translationally modified form (Bhagwat et al., 2021; Wild et al., 2019), can drive branch migration during meiosis. Other helicases (Mph1, Srs2 and Pif1) participate in certain aspects of meiotic recombination in budding yeast, but mutants lacking each of these helicases show normal or near-normal CO levels (Hunt et al., 2019; Sandhu et al., 2020; Vernekar et al., 2021).

In summary, the finding that branch migration is near-ubiquitous during meiotic CO formation prompts us to suggest that helicase-driven branch migration is a central feature of dHJ formation. In the future, it will be of interest to test this model, by identifying mutants and DNA sequence features that affect branch migration and determining their impact on meiotic recombination.

It should be noted that that the distances between final dHJ positions inferred from the location of strand switches in COs (median of 0.5-0.7 kb, up to ∼4kb; Figure 5; Figure S5A) are mostly greater than inter-junction distances (∼250 nt) in meiotic dHJs detected by electron microscopy (Bell and Byers, 1983; Oh et al., 2007; Oh et al., 2008). This difference, which remains to be accounted for, may be caused junction movement during DNA isolation and/or during EM sample preparation, or may reflect the possibility that, if the two HJs are not resolved at the same time, one junction can move after the other is resolved.

### Crossovers frequently undergo resolution-associated strand processing

COs also displayed frequent full conversion of multiple markers in heteroduplex (Figure 6). Since the strains used here are MMR-deficient, it is likely that this involves removal and resynthesis of DNA strands without regards to their heteroduplex content. Such processes include the repair of gaps formed by two nearby DSBs (Johnson et al., 2019; Prieler et al., 2021), “short patch” repair (Coic et al., 2000; Crown et al., 2014; Fleck et al., 1999) initiating at spontaneous DNA lesions, and strand removal and resynthesis at nicks formed by HJ resolution (Cannavo et al., 2020; Kulkarni et al., 2020; Marsolier-Kergoat et al., 2018). Both gap-repair and short-patch repair should occur at similar levels in NCOs and COs, but the events we observed were highly skewed toward COs, making it likely that they involve dHJ resolution by the MutL*γ* protein complex. Consistent with this, CO-associated full conversions of 2 or more markers are strongly reduced in *mlh1*Δ, *mlh3*Δ and *exo1*Δ mutants (Marsolier-Kergoat et al., 2018; see also Getz et al., 2008).

We envision three ways this might occur. Strand removal might result from multiple rounds of PCNA-directed strand-specific nicking by Mlh1-Mlh3 (Kulkarni et al., 2020), by synthesis and strand-displacement by DNA polymerase recruited to resolution sites (“nick-translation”; Marsolier-Kergoat et al., 2018), or by the activity of ExoI, which is associated with MutL*γ*. It will be of interest, in the future, to determine the activities responsible for the resolution-associated strand processing.

Recent studies suggest that a substantial fraction (∼20%) of DSBs involve multiple cuts; these form gaps whose repair could result in 6:2 marker segregation (Johnson et al., 2019; Prieler et al., 2021). Consistent with this, we find that 11% of COs and 12% of NCOs contain full conversions in or adjacent to the ∼400 nt DSB region (File S1.1). These are a minor fraction (∼11%) of total full conversions among COs, indicating that most of the 6:2 marker segregations we observed are not due to gap-filling.

### Concluding remarks

We have shown that the NCOs and COs that form during budding yeast meiosis display similar patterns of break repair-associated heteroduplex, while COs, and not NCOs, frequently display hallmarks of HJ branch migration and resolution-associated strand processing. Based on these findings, we have proposed the disassembly/migration-annealing model for meiotic recombination, where most hybrid DNA in both COs and NCOs is formed by processes akin to SDSA, and where branch migration is an integral step in CO formation. This model can account for many features of meiotic recombination in budding yeast, and can also account for features of previous genetic studies in fungi (Fincham, 1974; Hamza et al., 1987; Rossignol et al., 1984; Theivendirarajah and Whitehouse, 1983). The extent to which it accurately describes events in other organisms remains to be determined. While data from other organisms are limited, studies of mouse meiosis indicate differences with budding yeast, including substantially shorter total NMS tract lengths, especially among NCOs, different NMS tract spectra in NCOs and COs, and a notable deficit of CO products with heteroduplex on both sides of the initiating DSB (Cole et al., 2014; Cole et al., 2010; Peterson et al., 2020). Despite these differences, features of mammalian data, including COs and NCO tracts remote from the initiating DSB (Cole et al., 2014; Cole et al., 2010; Peterson et al., 2020) are reminiscent of the outcomes of template switching and branch migration seen in budding yeast. It will be of considerable interest, as more high-resolution data become available, to compare parental strand contributions to meiotic recombinants in different organisms and to determine which recombination processes are shared, and which are different.

## MATERIAL AND METHODS

### Yeast strains

All strains are SK1 derivatives (Kane and Roth, 1974); genotypes are listed in Supplementary Table 1.16. Drug resistance-marked deletions and insertions were created as described (Longtine et al., 1998). The previously described *his4::URA3-ARG4* hotspot (Parent 1 in Figure 2A; Jessop et al., 2005) was modified to form *his4::URA3-ARG4-SNPs* (Parent 2 in Figure 1A) using synthetic DNA fragments and Gibson assembly. Sixty percent of the markers are G to C changes (File S1.2), and all preserve *URA3* and *ARG4* coding information. Sequences contained in the insert locus were deleted from the endogenous *URA3* and *ARG4* loci. Ochre mutations (*ura3-K54oc* and *arg4-E93oc*) were created by site-directed-mutagenesis. All constructs were confirmed by sequencing.

### Sporulation, tetrad dissection and Southern blots

Sporulation and Southern blots for molecular CO scoring were as described (Kaur et al., 2018), with the addition of quick mating to avoid accumulation of recessive lethal mutations in *msh2*Δ diploids. For liquid sporulation, 5 colonies (56h at 30°C on YPD agar plates; 2% glucose, 2% Bacto Peptone, 1% Bacto Yeast Extract, 2% Bacto agar) each from *MAT***a** and *MAT*α parents were mixed in 20 µL of YPD broth, spotted on a YPD plate and incubated at 30°C for 7h. The patch was resuspended in 4 mL YPD broth containing 300µg/mL hygromycin (Invitrogen) and aerated at 30°C for 12 h, cells were pelleted and resuspended in 4 mL YPD broth containing 100 µg/mL nourseothricin sulfate (Neta Scientific) and aerated at 30°C for 12 h. This culture was then used to inoculate overnight SPS presporulation cultures as described (Kaur et al., 2018). DNA preparation was as described (Ahuja and Börner, 2011) without crosslinking. Probes are listed in File S1.17. Most tetrads (n=124) used for sequencing were from a 24 hr liquid sporulation culture and were dissected onto YPD agar plates. The rest were from quick matings as above, but cells were directly patched from YPD agar mating plates to 1% potassium acetate, 2% Bacto agar plates and were dissected after at least 2 days incubation at 30°C. For analysis of strains heterozygous for *ura3-K54oc* and *arg4-E93oc* alleles, tetrads were dissected onto YPD plates but with 4% glucose and 2.2% Bacto agar, which allowed more reliable detection of sectored colonies after replica plating.

### DNA preparation and sequencing

Entire spore colonies were inoculated into wells of deep-well 96 well plates (USA Scientific,1896-2110) with 500 µL of YPD broth/well and incubated overnight at 30° with aeration. 400 µl of each culture was used to prepare DNA, and the remainder was archived at -80°C. Cells were centrifuged at 3220xG for 2 min, the supernatant was removed, and cells were resuspended in 0.2 ml of 10mM NaPO_4_ pH 7.2, 10 mM EDTA, 0.1% ß-mercaptoethanol, 100 µg/ml Zymolyase T100 (MP Biomedicals). Cells were incubated 30 min at 37°C with tap-mixing every 5 min. Spheroplasts were centrifuged 10 min, 300 × g, the supernatant was removed, and spheroplasts were resuspended in 180 µl ATL buffer (Qiagen) with 0.5 mg/mL proteinase-K (Roche) and incubated at 65°C for 30 min, followed by addition of 5 µl of 4 µg/mL DNase-free RNase (Roche). Samples were loaded onto 96-well Qiacube-HT® columns and DNA was purified using a Blood & Tissue kit (Qiagen). After quantification, 1ng of DNA was used to PCR amplify the entire 6.865 kb recombination interval using the following primers: ACGGCACCACTATAAACCCG and GTGGGCTAAAGAACGCGAAC, and Q5 DNA polymerase (New England Biolabs) as recommended by the manufacturer, except that an extension time of 1 min/kb was used to avoid partial synthesis and priming by partial synthesis products.

Most of the samples were sequenced on a MiSeq sequencer (Illumina) after barcoding with a Nextera-384 barcoding kit (Illumina), resulting in ∼20 million reads/run that were up to 300 bp in length, with a typical coverage of ∼300-400 unique (deduplicated) reads/nucleotide/sample. The remaining samples were amplified from spore colony DNA using barcodes from (Guo et al., 2015) attached the outer primers listed above, and the entire region was sequenced on a Pacbio RS II instrument.

For Illumina sequencing, individual spore DNA sequences were aligned to the recombination interval sequence (Li and Durbin, 2010) followed by extraction of read base composition at each marker position, initially using a custom script and later using the sam2tsv function of jvarkit (Lindenbaum, 2016). For reads with more than one base at a marker position (i.e from a colony with heteroduplex), strand phase was determined manually from overlapping but non-conflicting reads and SNP abundance plots; in cases where phase was ambiguous, interval DNA from a single colony struck from archived samples was sequenced, initially by Illumina and later by PacBio sequencing (File S1.3). PacBio reads were demultiplexed using the locate function of seqkit (Shen et al., 2016), sample reads were aligned using minimap2 (Li, 2018), base composition per nucleotide was determined using the sam2tsv function of the jvarkit as above, and phase was determined manually from individual full-length reads.

### Data deposition and analysis

Sequencing data is deposited at the Sequence Read Archive (https://www.ncbi.nlm.nih.gov/sra) under BioProject ID: PRJNA721091. Underlying data for all figures are in File S1, a Microsoft Excel workbook. References to File S1 in the text use the following format: File S1.x, where x is the worksheet number. Statistical and other data analysis was performed using Graphpad Prism and R (https://www.R-project.org/). Plots of individual tetrad genotypes are in File S2, and were constructed using the R packages tidyverse and ggplot2 (Wickham et al, 2019; https://doi.org/10.21105/joss.01686). For tetrad analysis, map distances and standard error were calculated as described (Perkins, 1949) as implemented at https://elizabethhousworth.com/StahlLabOnlineTools/.

## ACKNOWLEDGEMENTS

We thank Eric Alani, Julia Cooper, Neil Hunter, Alexander Kelly, Bertrand Llorente and Matthew Neale for discussions and comments on the manuscript, and the Center for Cancer Research Sequencing Facility for high-throughput sequencing. This work used resources at the NIH HPC Biowulf cluster (http://hpc.nih.gov) and is supported by the Intramural Research Program of the NIH through the Center for Cancer Research at the National Cancer Institute.

---

**Figure S1.**
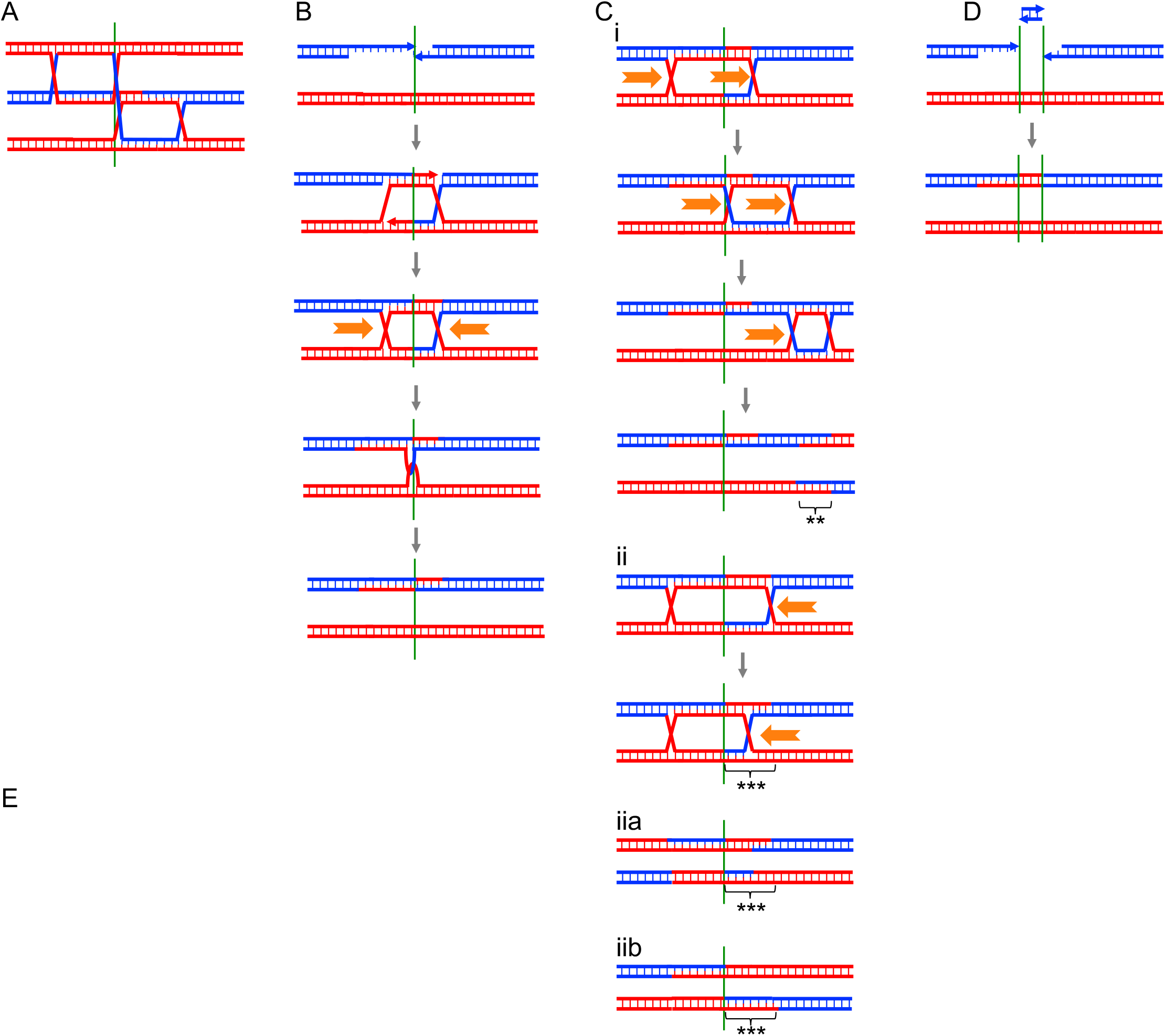
Other recombination mechanisms. (A) An example of multi-chromatid joint molecule in which a single DSB can engage more than one repair template and create hybrid DNA on three or more chromatids. (B) dHJ dissolution. Helicase-driven inward migration of the two Holliday junctions (orange arrows) followed by single-strand decatenation produces an NCO with hybrid DNA on both sides of the DSB. (C) Template switching during DSB end extension. End extension followed by D-loop disassembly provides a substrate for a second round of invasion and extension. If invasion alternates between the homolog and sister chromatid, a tract of mosaic heteroduplex (*) is formed. (D) Holliday junction branch migration. Holliday junction movement, by breaking old base-pairs and making new ones, can lead to dHJ migration to a new position. i) Movement of both junctions can result in crossovers and symmetrical hybrid DNA (**) that are separated from the DSB by parental DNA. Note that the hybrid DNA tract associated with the DSB is flanked by parental sequences in a noncrossover configuration. ii) Movement of one junction can produce hybrid DNA tracts that switch between parental molecules (***) at a point away from the DSB; resolution as a crossover produces either a hybrid DNA tract switch remote from the DSB (iia) or a canonical DSBR product (iib). (E) Multiple DSBs on a single chromatid produce a double-strand gap, which when repaired contains a patch of full conversion (6:2 segregation) in between the two DSBs.

**Figure S2.**
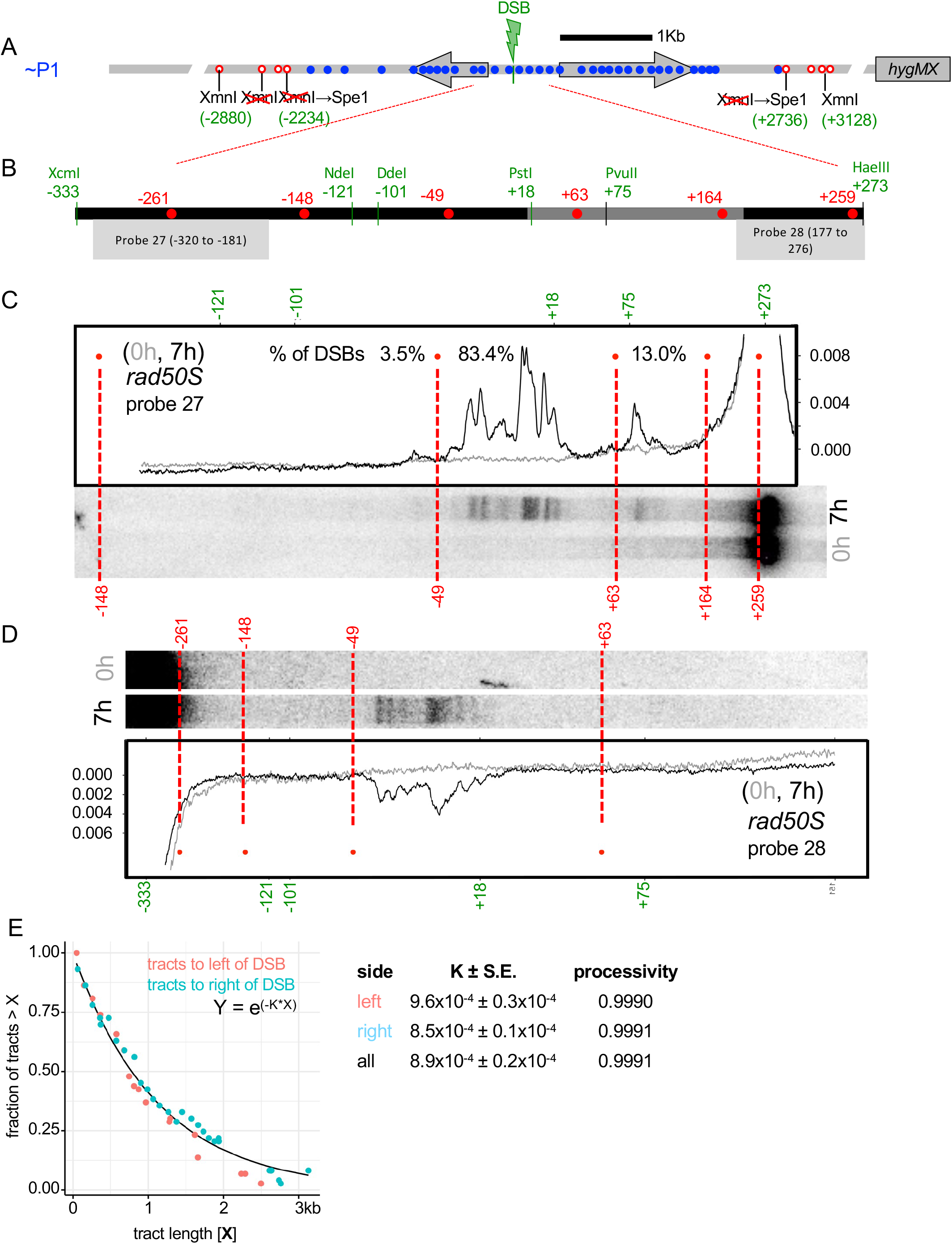
(A) Illustration of parent ∼P1, which lacks SNPs between nt -2236 and +2736, but contains flanking XmnI site polymorphisms to allow for scoring recombination on Southern blots. (B) Hotspot region (−333 to +273) used for high resolution DSB analysis. Green–restriction sites used for size standards; red–marker positions. (C, D) Blots of polyacrylamide gels with rad50S samples digested with XcmI and HaeIII and probed with probes 27 and 28, respectively, with plots of lane signal intensity. (E) Gene conversion tract lengths were fitted to an exponential decay curve. Processivity— fraction of tracts ≥ n nt that are ≥ n+1 nt, calculated as described (de Massy, 2003).

**Figure S3.**
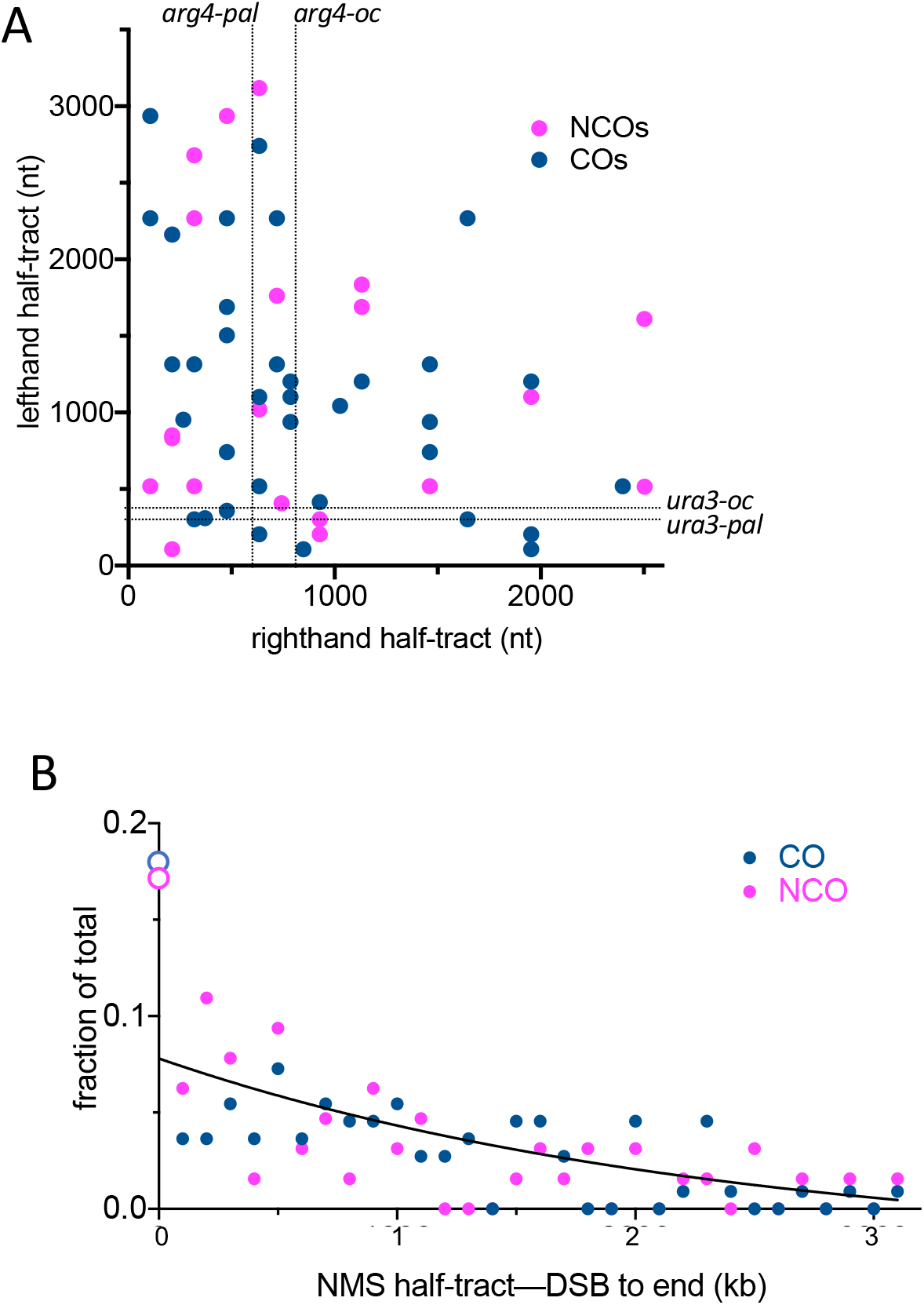
(A) Independent end extension in two-sided events. Half-tract lengths in two-sided COs (blue) and NCOs (magenta) are plotted. Left-hand and right-hand events are poorly correlated (Spearman rank-order correlation, p=0.09). Dotted lines denote location of arg4 and ura3 loss of function mutations used in conventional tetrad analysis in this study (Figure 2C and Supplementary Table 1.5; *arg4-oc* and *ura3-oc*) and in Jessop et al. (2005; *arg4-pal* and *ura3-pal*). Two-sided events detected by SNP analysis to the left of or below these lines would have been scored as one-sided had conventional marker analysis been used. (B) Non-cumulative distribution of NMS half-tract lengths, partitioned in 0.1 kb bins. Black line¬¬– exponential decay curve fitted to non-zero values. Open-circles—”invisible” half-tracts (<100nt), assuming that one-sided events contain a second “invisible” half-tract.

**Figure S4.**
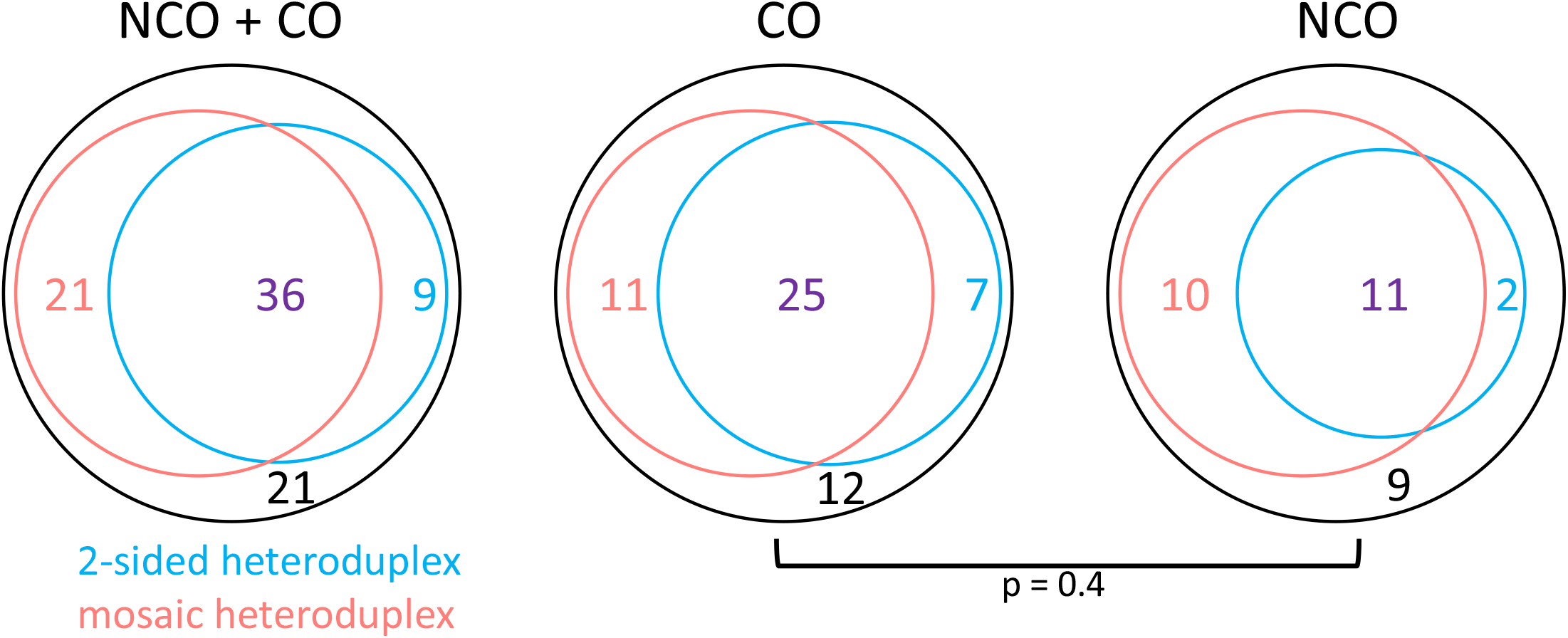
Multiple end invasions are common in both COs and NCOs. Venn diagrams showing number of events with 2-sided heteroduplex (both DSB ends invade), mosiaic heteroduplex (template switching), or both. Five crossover tetrads were too complex to score unambiguously. The distributions of events in NCOs and COs are not significantly different (chi-square test).

**Figure S5.**
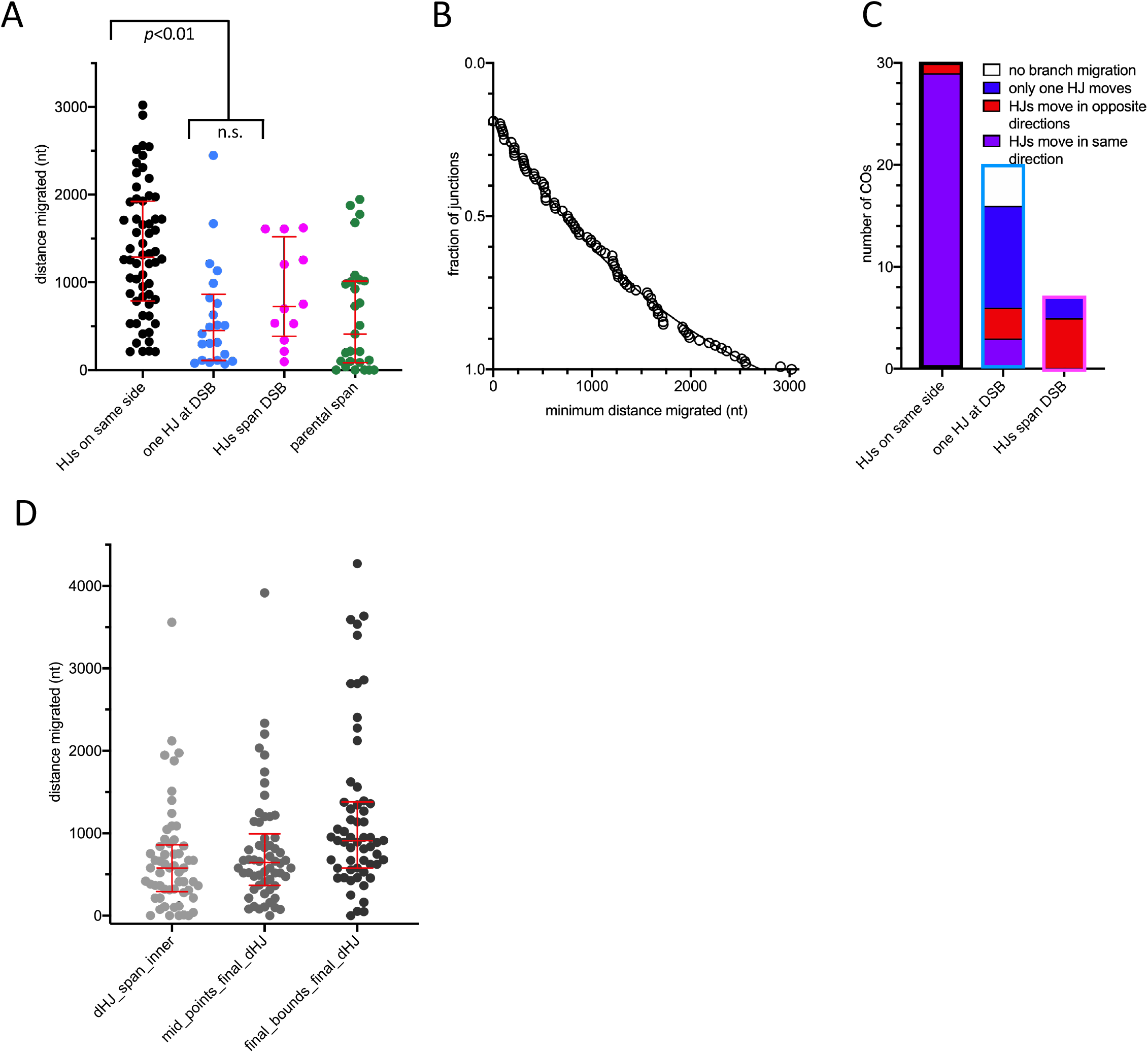
Properties of branch migration. A) Branch migration distances for each of the two Holliday junctions in COs, calculated by subtracting inferred initial junction positions from inferred final junction positions (see Figure 5A for examples). Black—final positions of both junctions are on one side of the DSB; turquoise— one junction is not separated from the DSB; pink—the two junctions are on opposite sides of the DSB (pink); green—extent of intervening parental sequences separating initial and final dHJ positions (see Figure 5A). Red bars denote median and quartiles. p values are from Mann Whitney rank tests. (B) rank order plot of branch migration distances, including 10 junctions where migration could not be detected. The curve is an exponential decay with a half-distance of 1kb, r2 = 0.99. (C) Types of branch migration for each class of crossovers. (D) Distance between final Holliday junctions; junction positions calculated relative to the two markers flanking the junction as follows: minimum—at the inner pair of flanking markers; midpoint—halfway between the flanking markers; maximum—at the outer pair of flanking markers. Red bars denote median and 1st and 3rd quartiles.

**Figure S6.**
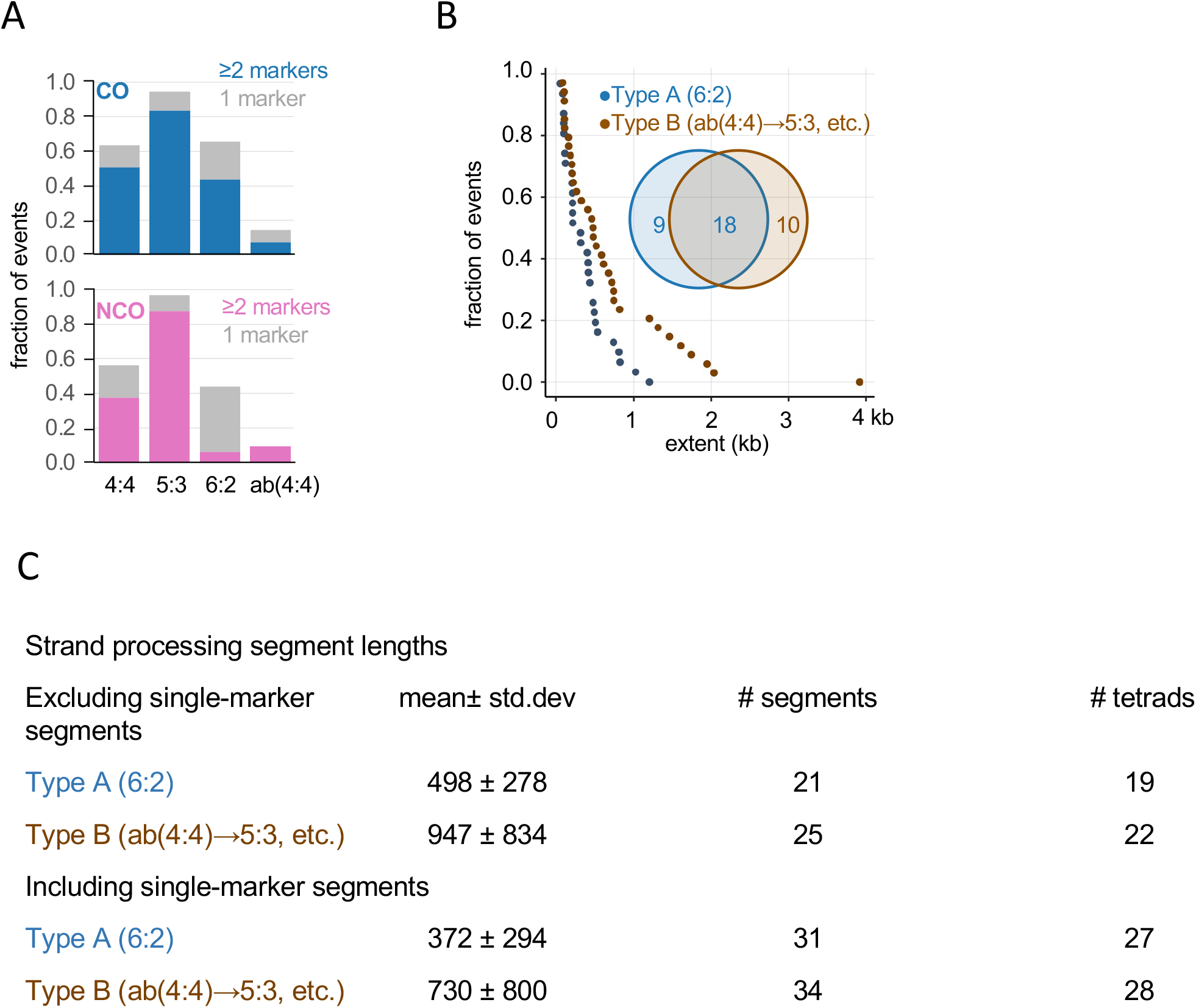
Strand processing signatures among crossovers. A) abundance of various marker segregation types among tetrads in the crossover and the noncrossover categories. (B) Rank order plots of strand processing tract lengths, including single-marker tracts. Median tract lengths: type A—316 nt; type B— 478 nt. Inset—Venn diagram showing number of tetrads with Type A, Type B, and both types of processing if single marker events are included. (C) Other features of strand processing segments.

**Figure S7.**
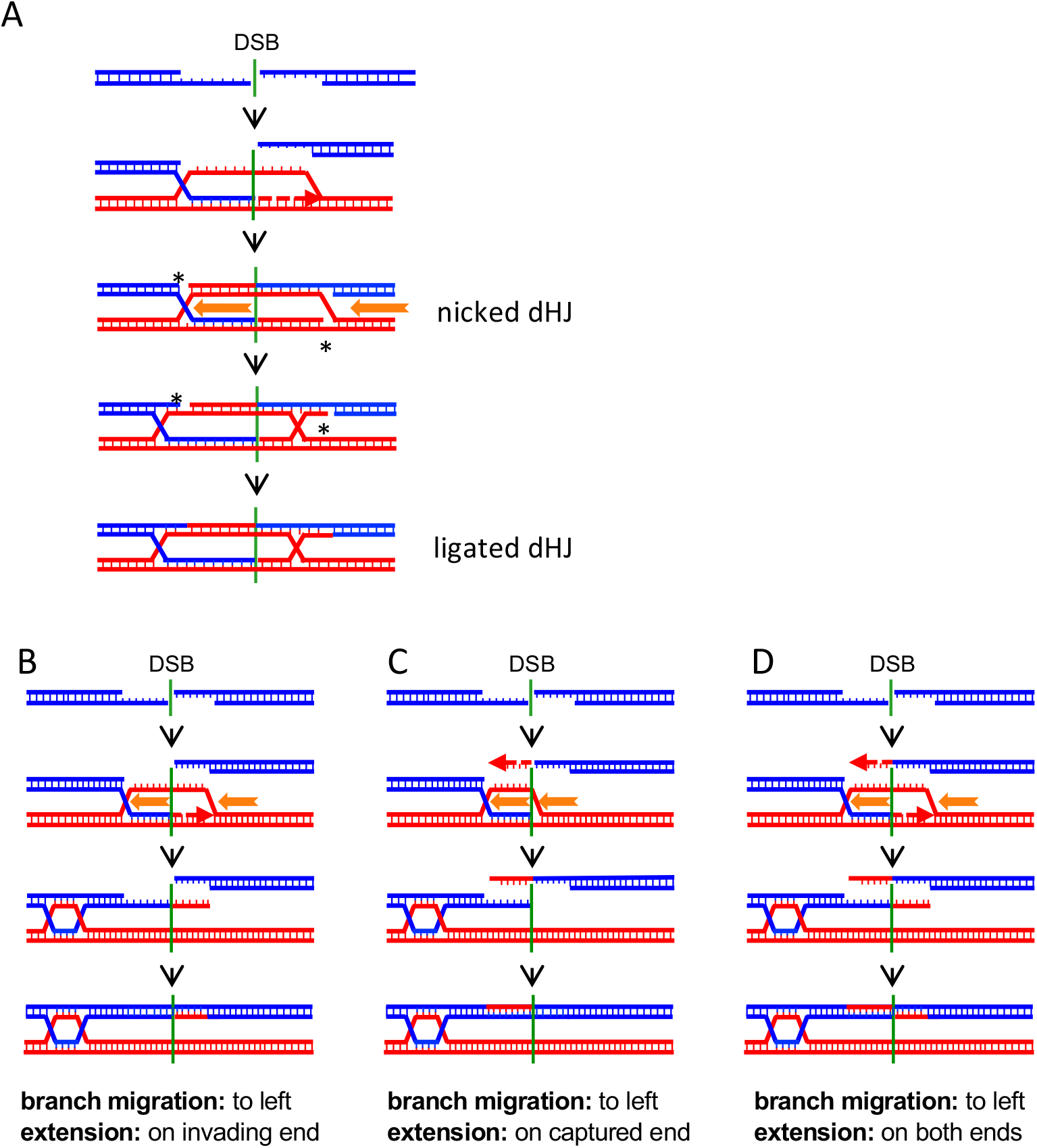
Different branch migration and annealing regimes. A) In the canonical DSBR model, where annealing immediately follows invasion and extension (Szostak et al., 1983), branch migration (orange arrows) for a short distance is required to move strand discontinuities at each branch (asterisks) into duplex DNA, where they can be ligated. B) Branch migration as in Figure 7. Synthesis from the invading end, followed by branch migration and annealing with an unextended DSB end, produces a crossover and hybrid DNA on opposite sides of the DSB. C) Invasion without synthesis, followed by branch migration and annealing with an extended DSB end, produces a CO and hybrid DNA on the same side of the DSB. D) Synthesis from the invading end, followed by branch migration and annealing with an extended DSB end, produces a CO with hybrid DNA on both sides of the DSB.

