## Supplemental File S2 for "Repeated strand invasion and extensive branch migration are hallmarks of meiotic recombination": File S2.html

Tetrad plots:


### Tetrad plots:

---

#### Tetrad 6

#### Tetrad 14

#### Tetrad 17

#### Tetrad 22

#### Tetrad 26

#### Tetrad 34

#### Tetrad 46

#### Tetrad 62

#### Tetrad 92

#### Tetrad 96

#### Tetrad 99

#### Tetrad 102

#### Tetrad 104

#### Tetrad 108

#### Tetrad 110

#### Tetrad 119

#### Tetrad 121

#### Tetrad 141

#### Tetrad 157

#### Tetrad 167

#### Tetrad 172

#### Tetrad 175

#### Tetrad 176

#### Tetrad 178

#### Tetrad 179

#### Tetrad 182

#### Tetrad 188

#### Tetrad 198

#### Tetrad 207

#### Tetrad 208

#### Tetrad 213

#### Tetrad 220

#### Tetrad 231

#### Tetrad 232

#### Tetrad 240

#### Tetrad 241

#### Tetrad 254

#### Tetrad 261

#### Tetrad 267

#### Tetrad 272

#### Tetrad 274

#### Tetrad 278

#### Tetrad 285

#### Tetrad 288

#### Tetrad 290

#### Tetrad 295

#### Tetrad 298

#### Tetrad 315

#### Tetrad 323

#### Tetrad 347

#### Tetrad 349

#### Tetrad 362

#### Tetrad 382

#### Tetrad 404

#### Tetrad 410

#### Tetrad 3

#### Tetrad 16

#### Tetrad 25

#### Tetrad 44

#### Tetrad 65

#### Tetrad 67

#### Tetrad 77

#### Tetrad 78

#### Tetrad 95

#### Tetrad 152

#### Tetrad 156

#### Tetrad 161

#### Tetrad 177

#### Tetrad 200

#### Tetrad 205

#### Tetrad 221

#### Tetrad 226

#### Tetrad 229

#### Tetrad 233

#### Tetrad 244

#### Tetrad 251

#### Tetrad 259

#### Tetrad 265

#### Tetrad 286

#### Tetrad 299

#### Tetrad 303

#### Tetrad 316

#### Tetrad 317

#### Tetrad 369

#### Tetrad 374

#### Tetrad 391

#### Tetrad 405

#### Tetrad 15

#### Tetrad 18

#### Tetrad 55

#### Tetrad 58

#### Tetrad 63

#### Tetrad 90

#### Tetrad 91

#### Tetrad 98

#### Tetrad 105

#### Tetrad 120

#### Tetrad 151

#### Tetrad 230

#### Tetrad 260

#### Tetrad 264

#### Tetrad 273

#### Tetrad 277

#### Tetrad 309

#### Tetrad 314

#### Tetrad 320

#### Tetrad 326

#### Tetrad 343

#### Tetrad 393

---

##### Legend
